# Sea anemone Frizzled receptors play partially redundant roles in the oral-aboral axis patterning

**DOI:** 10.1101/2022.03.15.484449

**Authors:** Isabell Niedermoser, Tatiana Lebedeva, Grigory Genikhovich

## Abstract

Canonical Wnt (cWnt) signaling is involved in a plethora of basic developmental processes such as endomesoderm specification, gastrulation and patterning the main body axis. To activate the signal, Wnt ligands form complexes with LRP5/6 and Frizzled receptors, which leads to nuclear translocation of β-catenin and transcriptional response. In Bilateria, the expression of different Frizzled genes is often partially overlapping, and their functions are known to be redundant in several developmental contexts. Here we demonstrate that all four Frizzled receptors take part in the cWnt-mediated oral-aboral axis patterning in the cnidarian *Nematostella vectensis* but show partially redundant functions. However, we do not see evidence for their involvement in the specification of the endoderm – an earlier event likely relying on maternal, intracellular β-catenin signaling components. Finally, we demonstrate that the main Wnt ligands crucial for the early oral-aboral patterning are Wnt3 and Wnt4. Comparison of our data to the knowledge from other models suggests that distinct but overlapping expression domains and partial functional redundancy of cnidarian and bilaterian Frizzled genes may represent a shared ancestral trait.

## Introduction

Wnt ligands and their Frizzled (Fz) receptors are involved in multiple cellular signaling pathways, one of which leads to the nuclear accumulation of β-catenin and is termed the “canonical” Wnt/β-catenin pathway or the cWnt pathway (MacDonald and He, 2012; van Amerongen and Nusse, 2009). In the “cWnt-off” state, cytosolic β-catenin is continuously tagged for degradation by the “destruction complex” containing APC, Axin, CK1α and GSK3β (Grainger and Willert, 2018), ubiquitinated by β-TrCP and degraded by the proteasome (Aberle et al., 1997). In the “Wnt-on” state, a complex of Wnt, Fz and the co-receptor LRP5/6 forms at the membrane, which results in the sequestering of the destruction complex by Dishevelled, which, in turn, prevents tagging β-catenin for degradation (Gammons and Bienz, 2018; Willert et al., 1999). Non-tagged β-catenin accumulates in the cytosol and becomes translocated into the nucleus, where it displaces the transcriptional co-repressor Groucho, and interacts with TCF to activate target genes (Flack et al., 2017). In addition to the role in cWnt signaling, which is characterized by the involvement of LRP5/6 and the nuclear translocation of β-catenin, Wnt ligands and Fz receptors are the starting points of multiple “non-canonical” signaling pathways (Acebron and Niehrs, 2016; Croce and McClay, 2008; Garcia de Herreros and Dunach, 2019; Park et al., 2015; Semenov et al., 2007; Villarroel et al., 2020). In mammals, ten different Fz receptors comprising five families may demonstrate partially overlapping functions, and the effects of their individual or combined knockouts are usually attributed to a mixed action of the abnormal cWnt and non-canonical signaling (Fischer et al., 2007; Wang et al., 2016). Among the mammalian Fz receptors, only Fz4 appears to act exclusively in the cWnt pathway, while Fz3 and Fz6 seem to be exclusively involved in the Wnt/PCP pathway (Wang et al., 2016).

One of the ancestral roles of the cWnt pathway is to define the gastrulation site, as well as to pattern the main body axis in animals - a feature, which appears to be conserved across Metazoa. Localized expression of the Wnt signaling components along the main body axis has been documented in the earliest branching animal lineages such as ctenophores (Pang et al., 2010) and sponges (Adamska et al., 2010; Leininger et al., 2014). In Cnidaria, the bilaterian sister group, the role of the cWnt pathway in gastrulation and oral-aboral (OA) axis patterning has been confirmed by functional analyses (Kraus et al., 2016; Lebedeva et al., 2021; Leclère et al., 2016; Marlow et al., 2013; Momose et al., 2008; Momose and Houliston, 2007; Röttinger et al., 2012; Wikramanayake et al., 2003). Recently, we demonstrated that the regulatory logic of the β-catenin-dependent OA patterning in the sea anemone *Nematostella vectensis* and the posterior-anterior (PA) patterning of deuterostome Bilateria is highly similar, suggesting a common evolutionary origin of the OA and the PA axes (Darras et al., 2018; Darras et al., 2011; Kiecker and Niehrs, 2001; Lebedeva et al., 2021; Nordström et al., 2002). Although the way *Nematostella* interprets different intensities of β-catenin signal is largely understood (Kraus et al., 2016; Lebedeva et al., 2021), we still have very little idea about which Wnt ligands and which Fz receptors are involved in the cWnt-dependent axial patterning in this morphologically simple model organism. The complement of Wnt and Fz molecules in *Nematostella* is surprisingly large. It has representatives of 12 out of the 13 conserved bilaterian *Wnt* gene families, only lacking *Wnt9*, which has been lost in Cnidaria but is present in the earlier branching Ctenophora (Kusserow et al., 2005; Lee et al., 2006; Pang et al., 2010). *Nematostella* Wnt genes are expressed in staggered domains along the OA axis, with different Wnt sets transcribed in the ectoderm and in the endoderm (Kusserow et al., 2005; Lee et al., 2006). The *Nematostella* genome also harbors representatives of four out of five vertebrate Frizzled receptor families, *Fz1/2/7* (*Fz1* in the text below), *Fz4, Fz5/8* (*Fz5* in the text below), and *Fz9/10* (*Fz10* in the text below), and lacking only *Fz3/6*, which appears to be chordate-specific (Bastin et al., 2015; Schenkelaars et al., 2015).

In this study, we asked, which of the four Fz receptors and the many Wnt ligands are involved in the cWnt-dependent patterning of the oral-aboral axis in the *Nematostella* embryo. Since the involvement of LRP5/6 is the hallmark of the cWnt signaling, we reasoned that analyzing its loss-of-function phenotypes would tell us which parts of the OA patterning process are under cWnt control thus facilitating the interpretation of the Fz loss-of-function data. We show that the knockdown of LRP5/6 suppresses the expression of the β-catenin-dependent oral and midbody genes and expands aboral molecular identity without affecting endoderm specification. This results in a loss of the oral structures after gastrulation and a global expansion of the aboral/anterior molecular identity – a typical β-catenin loss-of-function phenotype. Individual knockdowns of the three orally expressed *Fz* do not affect oral marker gene expression. In contrast, dual-and triple-knockdowns of all possible *Fz* combinations partially phenocopy the *LRP5/6* knockdown, while quadruple *Fz* knockdown replicates it at the molecular and morphological level. These data suggest partial redundancy of the Fz receptors and involvement of all the *Nematostella* Fz receptors in cWnt-dependent OA patterning. We also demonstrate that Wnt3 and Wnt4 are the key Wnt ligands mediating OA patterning during early development.

## Results

### Normal expression of the *Fz* and *LRP5/6* genes in *Nematostella*

We cloned *Fz, LRP5/6* and *LRP4/5/6-like* sequences from cDNA and analyzed their temporal and spatial expression dynamics in *Nematostella* embryos and larvae by interrogating the NvERTx RNA-Seq database (Warner et al., 2018) and by performing whole-mount in situ hybridization. Transcriptomics data show that two out of four *Fz* genes, *Fz1* and *Fz5*, and *LRP5/6* are abundant in the unfertilized egg, and their expression is maintained at an approximately constant level. In contrast, the other two *Fz* genes, *Fz4* and *Fz10*, are zygotic and become activated around 8 hours post-fertilization (hpf) (Suppl. Fig.1A). The LRP4/5/6-like (Suppl. Fig. 1B) is a weakly expressed gene, which starts to be upregulated around 48 hpf, and its expression becomes confined to the forming apical organ (Suppl. Fig. 1C). Unlike the intracellular domain of *Nematostella* LRP5/6, the intracellular domain of *Nematostella* LRP4/5/6-like does not contain the PPPS/TP sites shown to be phosphorylated by GSK3 and act as Axin binding sites in bilaterian models (Tamai et al., 2004). Thus, we reasoned that LRP4/5/6-like is unlikely to be involved in cWnt signaling (at least not before 48 hpf) and did not consider it further.

ISH analysis of the *Fz1, Fz5*, and *LRP5/6* (Fig. 1) show the initially ubiquitous distribution of the mRNA prior to 10 hpf, which is then followed by the formation of a clearing in the expression likely corresponding to the future preendodermal plate. At the same time, *Fz4* and *Fz10* expression starts to be detectable. As the development progresses, a second clearing in the *Fz10* expression domain appears on the putative aboral end, while *Fz5* expression becomes most prominent aborally (see also Lebedeva et al., 2021). At the onset of gastrulation, *Fz1, Fz4* and *LRP5/6* are weakly expressed ubiquitously; additionally, LRP5/6 is becoming ever more prominent in the aboral ectodermal domain. *Fz10* is expressed in the oral and midbody ectoderm, but it also starts to be strongly expressed in the invaginating endodermal plate. At late gastrula, a narrow clearing in the *Fz1* expression starts to appear between the midbody ectoderm and the aboral ectoderm, and *Fz5* acquires an additional expression domain in the aboral endoderm. During planula development, *Fz1* expression forms an oral-to-aboral gradient with the maximum in the oral ectoderm and oral endoderm, however, *Fz1* transcript is also detectable in the apical organ. Apart from the apical organ expression, aboral ectoderm appears to be free of *Fz1* mRNA. *Fz4* transcription forms a shallow oral-to-aboral gradient in both germ layers, however, in contrast to *Fz1, Fz4* is not expressed in the apical organ. *Fz5* is expressed in an aboral-to-oral gradient in both cell layers with the ectodermal expression fading out at the aboral/midbody boundary. Apical organ cells express *Fz5* particularly strongly, and there it is co-expressed with *Fz1*. Strong *Fz10* expression is detectable in the pharyngeal, oral and midbody ectoderm. Additionally, *Fz10* forms an oral-to-aboral gradient of expression in the endoderm. Finally, LRP5/6 is expressed ubiquitously, however, apical organ cells appear to produce much more *LRP5/6*, and a shallow aboral-to-oral gradient appears to exist in the endoderm (Fig. 1).

**Figure 1.**
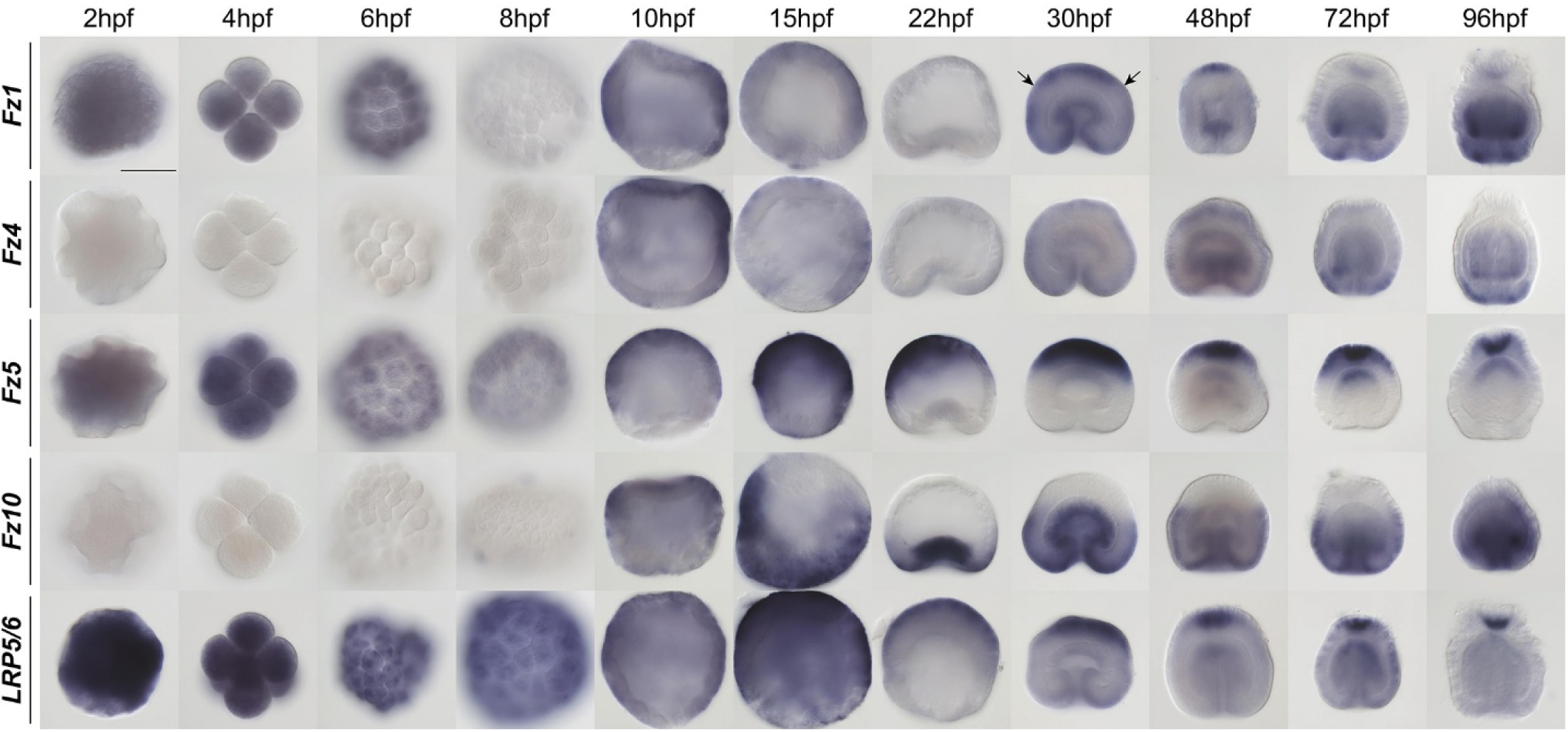
Normal developmental expression of the *Fz* genes and *LRP5/6*. From 10 hpf on, the animal/oral pole of the embryo is pointing down. Arrows point at the clearing in the *Fz1* expression developing at the midbody/aboral border. Scale bar 100 μm.

### LRP5/6 knockdown

To assess the role of *LRP5/6* we performed shRNA-mediated knockdowns (KD, Suppl. Fig. 2A-B) and characterized their effect on marker gene expression. We used *Brachyury* (*Bra*), *Wnt2* and *Six3/6* as markers of the oral, midbody and aboral domains respectively (Lebedeva et al., 2021), *Axin* as a β-catenin signaling target gene with broader expression (Kraus et al., 2016; Lebedeva et al., 2021), as well as several additional markers for specific areas in the embryo. Notably, the midbody marker *Wnt2* is also positively regulated by β-catenin signaling but it gets suppressed orally by Bra (Lebedeva et al., 2021). At the late gastrula stage (30 hpf), *LRP5/6* RNAi resulted in a strong suppression of the oral markers *Bra, FoxA* and *FoxB*, as well as *Axin* (Fig. 2, Suppl. Fig. 3). *Wnt2* was reduced and only detectable in the oral domain, while *Six3/6* strongly expanded orally and acquired an additional area of expression in the pharyngeal ectoderm (Fig. 2). The suppression of the oral ectodermal and the expansion of the aboral ectodermal domain signature into the oral ectodermal territory persisted into later developmental stages even though *LRP5/6* expression was re-established by 3 days post fertilization (dpf) (Suppl. Fig. 4). Despite normal gastrulation, oral and pharyngeal structures were later lost, and by 4 dpf all LRP5/6 RNAi embryos resembled diploblastic spheres (Fig. 3A). In summary, LRP5/6 RNAi phenocopied the outcome of dominant-negative *Tcf* (*dnTcf*) mRNA overexpression (Röttinger et al., 2012), and was strikingly similar to the effect of the combined KD of *Bra, Lmx, FoxA*, and *FoxB* - the four β-catenin-dependent transcription factors determining the oral molecular identity of the embryo (Lebedeva et al., 2021). LRP5/6 RNAi resulted in a typical β-catenin loss-of-function phenotype, apart from the obvious fact that the embryos gastrulated normally, which was also the case in *dnTcf* mRNA-injected embryos (Röttinger et al., 2012) but, curiously, not in β-catenin morphants (Leclère et al., 2016), or in embryos subjected to shRNA-mediated β-catenin RNAi (Karabulut et al., 2019). Unlike *LRP5/6* RNAi and *dnTcf* overexpression, β-catenin morpholino injection resulted in a complete suppression of the oral, midbody, and aboral ectoderm markers and in a ubiquitous upregulation of the endodermal marker *SnailA* (Leclère et al., 2016). In contrast, pharmacological activation of β-catenin signaling with azakenpaullone (AZK) starting at fertilization, also blocked gastrulation, however, in this case, *SnailA* expression was abolished, and oral ectoderm markers were expressed ubiquitously instead (Leclère et al. 2016, see also Fig. 4A). Curiously, endodermal marker expression, as well as the gastrulation process, was not affected by AZK treatment if the treatment started after 6 hpf (Fig 4A), which corresponds to the reported time of the activation of the zygotic genome (Helm et al., 2013). This suggests that endoderm specification probably relies on maternally deposited mRNA and proteins and occurs prior to 6 hpf, and that once specified, the endoderm becomes insensitive to modulations in the β-catenin signaling at least until late gastrula stage. Moreover, normal gastrulation and endodermal marker gene expression in shLRP5/6 embryos (Fig. 4B) raises the possibility that endoderm specification and the gastrulation movements, although obviously β-catenin-dependent, may not require Wnt/Fz/LRP5/6-mediated signalling. To address this in more detail, we first asked how soon the effect of LRP5/6 knockdown started to manifest itself after the RNAi. Despite clear *LRP5/6* suppression as early as 6 hpf, the effect of *LRP5/6* RNAi on the sensitive β-catenin signaling target *Bra* was not apparent at 10 hpf, and only became observable at late blastula (18 hpf) stage (Suppl. Fig. 5). Since this comparatively late manifestation of the LRP5/6 RNAi effect, rather than endoderm specification and invagination being Wnt/Fz/LRP5/6-independent, may be the reason for the difference between the morpholino-mediated β-catenin KD and the RNAi-mediated *LRP5/6* KD, we repeated *LRP5/6* KD using a translation-blocking morpholino (MO, Suppl. Fig. 2C). By 30 hpf (late gastrula stage in controls), LRP5/6MO injection resulted in a phenotype similar to that of LRP5/6 RNAi, although more pronounced: not only *Bra*, but also *Wnt2* expression was abolished, and *Six3/6* was expanded throughout the whole ectoderm. In contrast to LRP5/6 RNAi, gastrulation was delayed in the morphants; nevertheless, just like in RNAi, the specification of the *SnailA*-positive, *Six3/6*-negative pre-endodermal plate took place normally (Fig. 5A). By 48 hpf, the LRP5/6MO embryos remained arrested in gastrulation, demonstrating a miniature blastopore lip and a slightly submerged endoderm (Fig. 5B). By 4 dpf, LRP5/6 morphants displayed the same “bi-layered aboralized sphere” phenotypes as the LRP5/6 RNAi embryos (Fig. 3A, Fig. 5B). Both RNAi and MO-mediated knockdown clearly show that LRP5/6 is required for the cWnt-mediated patterning of the ectoderm in *Nematostella*. The conspicuous lack of endodermal mesenteries in the 4 dpf LRP5/6 RNAi and morphant embryos is a clear sign of the disrupted BMP signaling resulting in the loss of the second, “directive” body axis (Genikhovich et al., 2015; Leclère and Rentzsch, 2014). Previously, we demonstrated that β-catenin is required for the onset of the expression of *BMP2/4* and *Chordin* – the core components of the BMP signaling network in *Nematostella* (Genikhovich et al., 2015; Kirillova et al., 2018; Saina et al., 2009). Surprisingly, upon LRP5/6 RNAi, the directive axis is formed, but later lost, as evidenced by *Chordin* expression, which is initially normal and bilaterally symmetric at the gastrula stage, but disappears by 3 dpf (Suppl. Fig. 6). In summary, we conclude that *LRP5/6* is required for the β-catenin-dependent patterning of the ectoderm along the OA axis, and for the maintenance of the directive axis, but we do not find evidence of its involvement in the specification of the endoderm.

**Figure 2.**
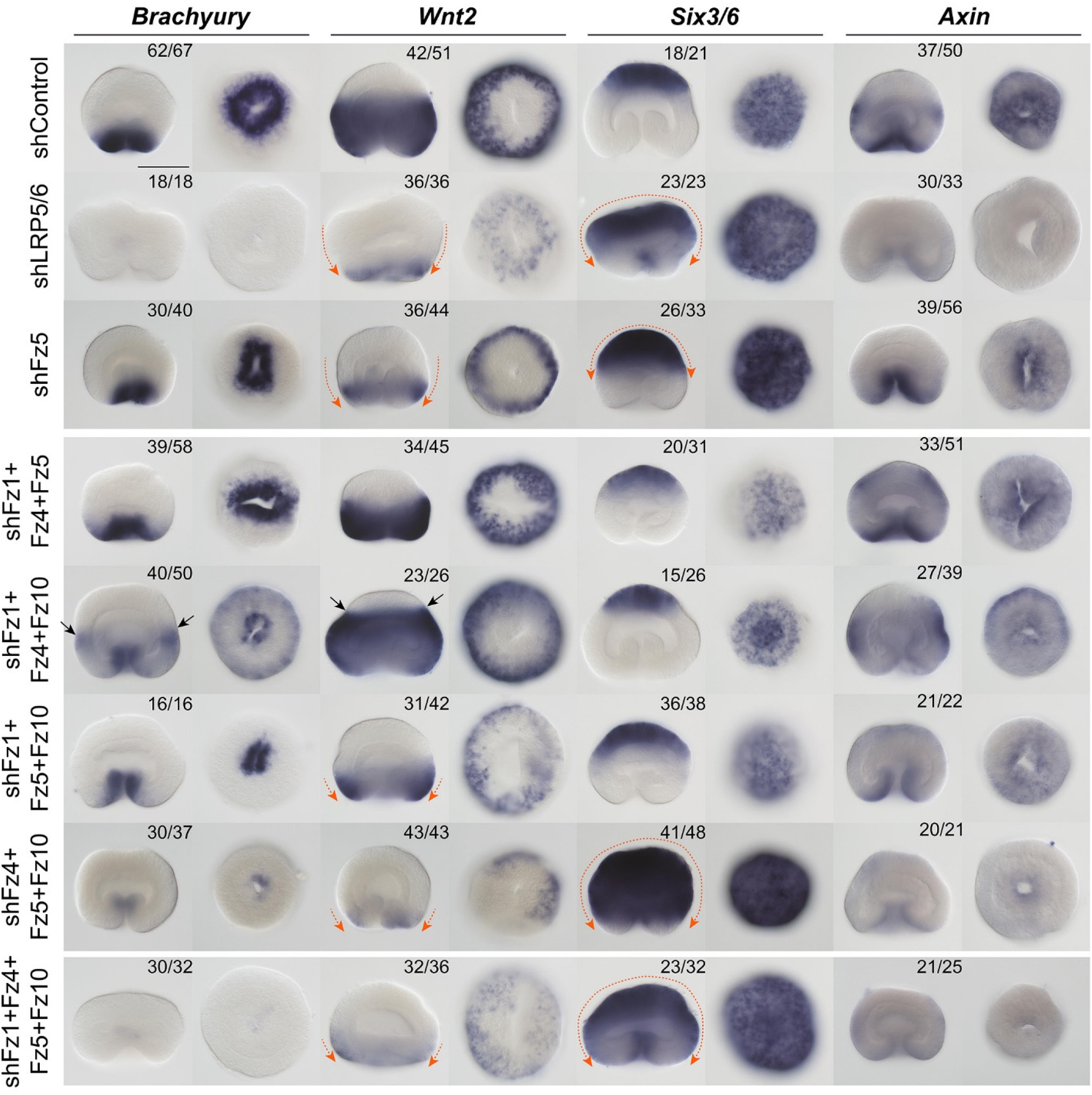
Effects of the RNAi-mediated KD of LRP5/6, Fz5, as well as triple and quadruple Fz combinations on the expression of the β-catenin-dependent markers of different axial domains. Orange arrows indicate the direction of the drastic expression shifts. Black arrows point at the ring of stronger *Bra* expression in the midbody and the aboral expansion of *Wnt2* domain suggesting an ectopic enhancement of the β-catenin signaling. The numbers in the top right corner show the fraction of the embryo demonstrating this phenotype. Scale bar 100 μm. For each gene, lateral views (oral end down) on the left, oral (aboral in case of *Six3/6*) views on the right.

**Figure 3.**
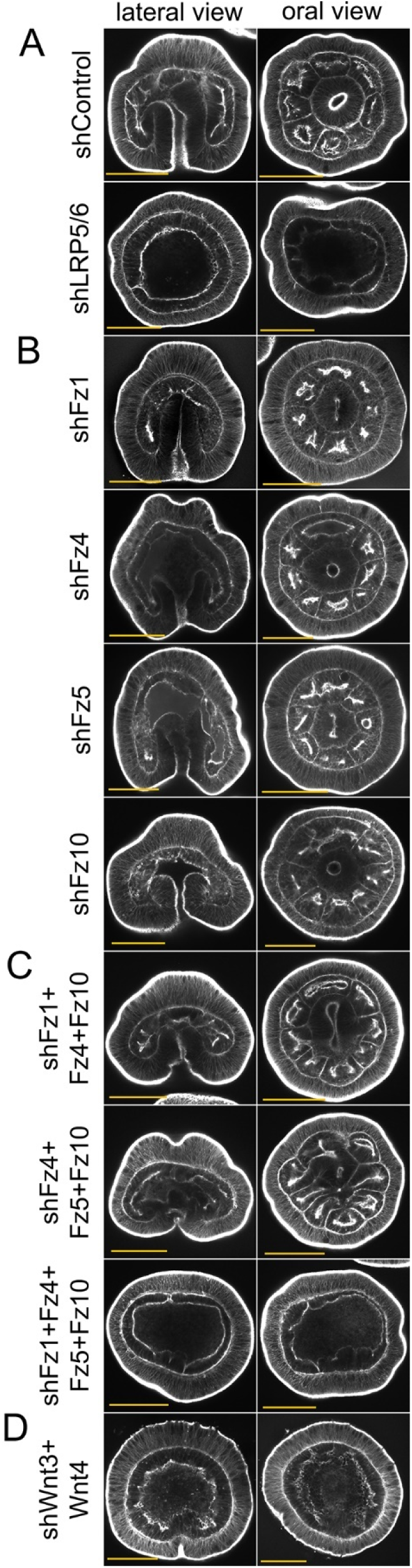
Effects of the RNAi-mediated KD of *LRP5/6* (A), individual *Fz* (B), triple and quadruple *Fz* KDs (C), and of the double KD of *Wnt3* and *Wnt4* (D) on the later development of the embryo. 4 dpf embryos are stained with phalloidin-AlexaFluor488. Scale bars 100 μm. On lateral views, oral end points down.

**Figure 4.**
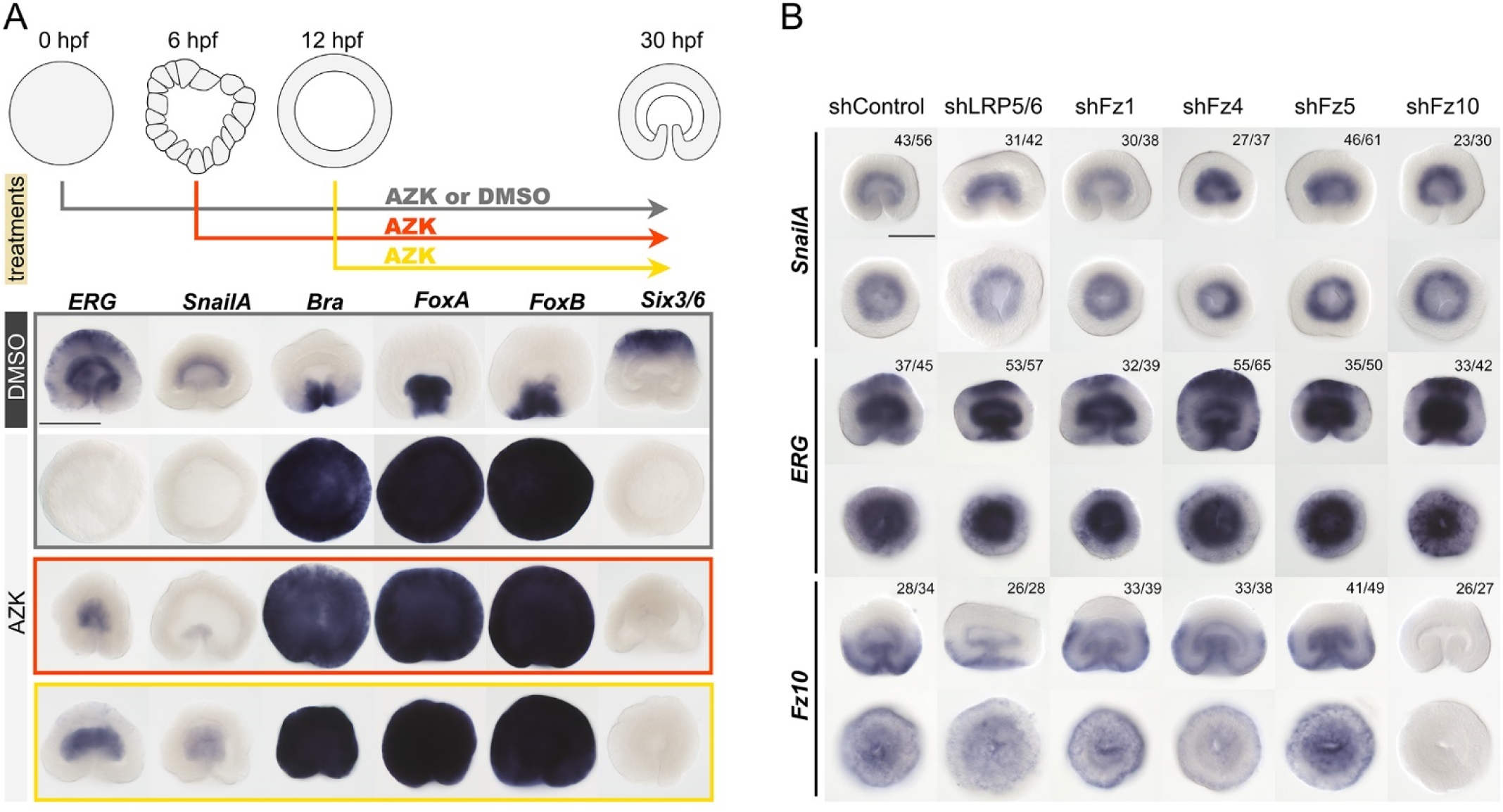
Endoderm specification is an early event, which does not seem to depend on Fz/LRP5/6. (A) Identification of the time of endoderm specification. Lateral views, oral end down. (B) Endodermal marker expression is not affected by the KDs of *LRP5/6* or individual knockdowns of *Fz*. The numbers in the top right corner show the fraction of the embryo showing this phenotype. For each gene, lateral views (oral end down) on the top, oral (aboral in case of *Six3/6*) views on the bottom. Scale bars 100 μm.

**Figure 5.**
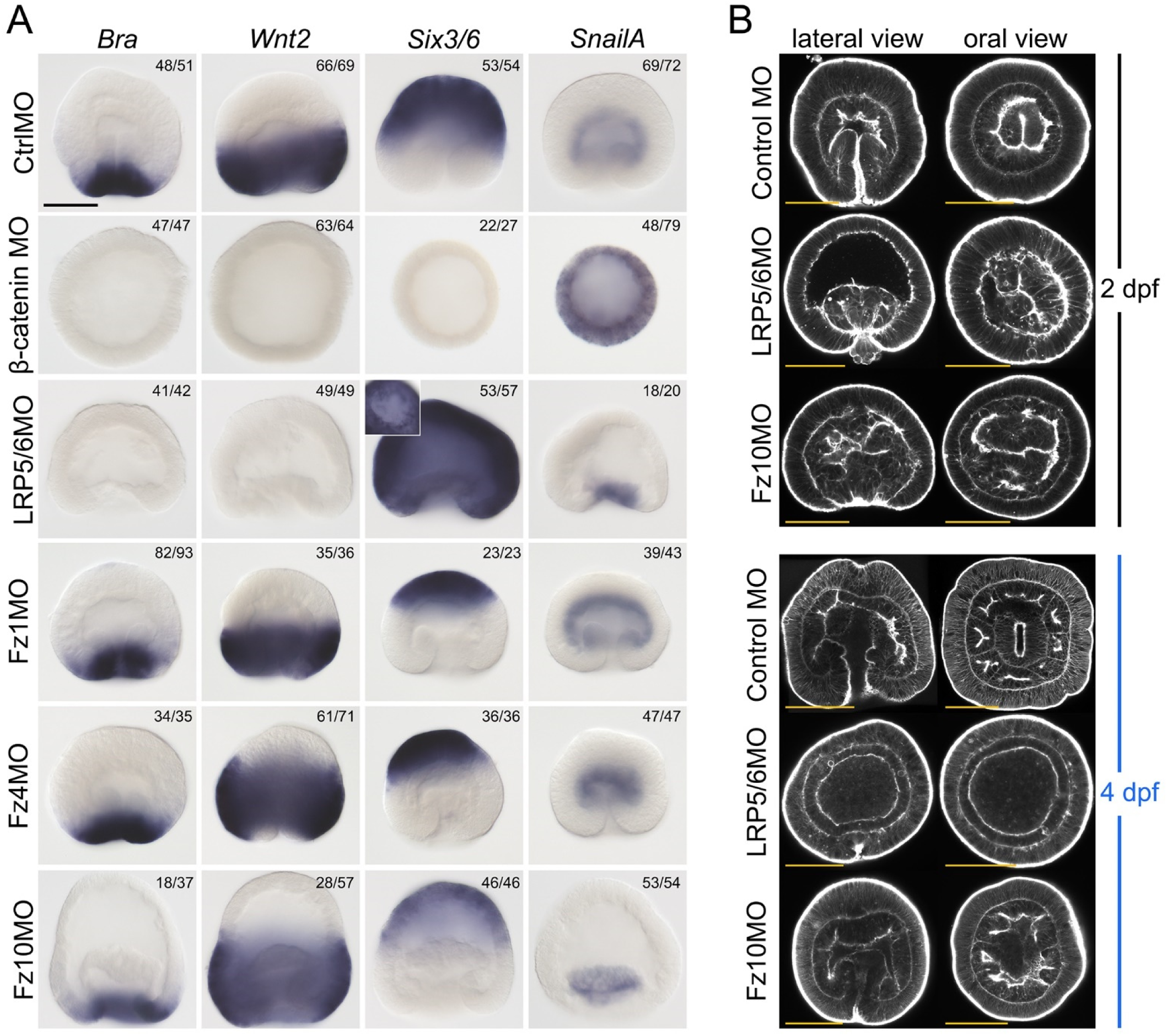
Effect of the morpholino-mediated KD of *LRP5/6* and orally expressed *Fz* genes on the early development of *Nematostella*. (A) Effect of the knockdowns on the expression of the markers of the distinct axial domains in the ectoderm, and on the endodermal marker *SnailA*. The numbers in the top right corner show the fraction of the embryo demonstrating this phenotype. Scale bar 100 μm. Lateral views, oral end down. The inset image of an oral view of the LRP5/6 morphant stained for *Six3/6* shows that the pre-endodermal plate does not express *Six3/6*. (B) Effects of the LRP5/6 and Fz10 morpholino KDs on the later development of the embryos. Scale bars 100 μm.

### Knockdown of Fz receptors

In contrast to *Fz5* RNAi, which reproduced the Fz5 morpholino knockdown phenotype published earlier (Leclère et al., 2016; this paper), individual RNAi of *Fz1, Fz4*, and *Fz10* did not result in changes in the *Bra, Wnt2, Six3/6* and *Axin* expression (Suppl. Fig. 7). The *Fz5* RNAi phenotype was similar to that of *LRP5/6* RNAi, with the aboral, *Six3/6*-expressing domain expanded, and the midbody *Wnt2*-expressing domain constricted towards the oral pole (Fig. 2). However, in contrast to *LRP5/6* RNAi, oral markers *Bra, FoxA*, and *FoxB* were not affected by *Fz5* RNAi, and only the midbody expression of *Axin* was suppressed, while oral expression was retained (Fig. 2, Suppl. Fig. 3). Endodermal expression of *SnailA, ERG*, and *Fz10* was also not affected by any of the *Fz* RNAi knockdowns except for *Fz10* expression, which, naturally, was abolished upon *Fz10* RNAi (Fig. 4B). Individual *Fz* RNAi did not lead to significant morphological defects apart from a slight gastrulation delay in *Fz1* and *Fz10* RNAi, and a previously reported slight shortening of the OA axis in *Fz5* RNAi (Leclère et al., 2016). By 4 dpf, the KD embryos developed eight normal mesenteries (Fig. 3B).

Surprisingly, these results contradicted the recently published *Fz1* and *Fz10* KD phenotypes (Wijesena et al., 2022). In this paper, the authors stated that overexpression of the dominant-negative form of *Fz1* (*dnFz1*) caused oral expansion of *Fz5*, suppression of *FoxA* in the blastopore lip, and the loss of the endodermal expression of *SnailA* and *Fz10* without interfering with the gastrulation process. In contrast, their Fz10 morpholino injection suppressed endoderm invagination without affecting *SnailA*. This latter result was somewhat surprising, since the disappearance of *Fz10* expression Wijesena et al. observed upon *Fz1* KD did not lead to a gastrulation failure. These results led the authors to conclude that Fz1 was controlling the cWnt-dependent specification of the endoderm, while Fz10 was regulating the non-canonical Wnt-dependent endoderm invagination (Wijesena et al., 2022).

Remembering the more pronounced effect of morpholino-mediated *LRP5/6* KD in comparison to RNAi, we repeated individual *Fz1, Fz4* and *Fz10* KD using the Fz1MO, Fz4MO and Fz10MO (Fig. 5, Suppl. Fig. 2C). Similar to the RNAi result, Fz1MO and Fz4MO injection did not lead to changes in the expression of the oral, midbody, aboral, and endodermal markers (Fig. 5A). In our hands, also the overexpression of *dnFz1-mCherry* mRNA did not cause any change in *Bra, Wnt2, Six3/6* and *SnailA* expression, just like the *Fz1* RNAi and Fz1MO KD (Suppl. Fig. 8). In contrast, Fz10 morpholino injection led to a delay in gastrulation without affecting *Bra, Wnt2, Six3/6* and *SnailA* expression (Fig. 5A). By 48 hpf, Fz10MO morphants completed invagination, although their endoderm still looked irregular and they had open blastopores (Fig. 5B). Their morphology mostly normalized by 96 hpf, with the only deviation being the lower number of mesenteries suggesting some developmental delay (Fig. 5B). Thus, it is likely that *Fz10* plays a role in regulating gastrulation; however, the similarity of the effect of Fz10 and LRP5/6 morpholino KD on the overall morphology of the gastrula raises the possibility that the gastrulation delay may be caused by the cWnt signaling-related defect. The proposed role of Fz10 in mediating non-canonical Wnt signaling cannot be excluded and has to be directly assessed in the future; however, we do not find clear support for the “Fz1 for cWnt and endoderm specification *vs*. Fz10 for non-canonical Wnt signaling and endoderm invagination” distinction proposed in Wijesena et al., 2022.

Since individual RNAi of the orally expressed *Fz* genes did not elicit an effect, we presumed that they might be partially or completely redundant at the gastrula stage, and performed simultaneous RNAi of all possible combinations of two, three, or four Frizzleds. Double *Fz* knockdowns showed effects on marker genes only if shFz5 was in the mix, and recapitulated the individual *Fz5* KD (Suppl. Fig. 7). In triple RNAi, the β-catenin loss-of-function phenotype similar to the *LRP5/6* RNAi started to emerge in some cases, most notably in the *Fz4*+*Fz5*+*Fz10* combination (Fig. 2). Simultaneous RNAi of *Fz1*+*Fz4*+*Fz10* resulted in a curious phenotype, which we are currently unable to explain: expression of *Bra* and *Axin* at the oral end of the gastrula became weaker, and a narrow ring of relatively strong *Bra* and a wider ring of strong *Axin* expression appeared in the midbody of the gastrula, suggesting stronger than usual β-catenin signaling in this area. This occurred concomitantly with the aboral expansion of the *Wnt2* domain and reduction of the *Six3/6* domain. *Wnt2* expression in this case was strongest in an area located between the Bra-expressing ring in the midbody and the diminished *Six3/6* expression domain (Fig. 2). In spite of the prominent effects at the gastrula stage, triple *Fz* RNAi embryos formed eight mesenteries by 4 dpf, although the mesenteries in the *shFz4*+*Fz5*+*Fz10* combination always looked somewhat irregular (Fig. 3C). Finally, quadruple RNAi of all four Fz receptors phenocopied LRP5/6 RNAi at the molecular as well as at the morphological level (Fig. 2, Fig. 3A,C). Taken together, we show that three orally expressed Fz receptors play a partially redundant function in the OA axis patterning of the *Nematostella* gastrula. The fact that only combined RNAi of all four Fz receptors phenocopies the LRP5/6 knockdown at the molecular and morphological level hints towards the involvement of all *Nematostella* Fz proteins in the LRP5/6-mediated cWnt signaling. We do not find evidence for the dependence of the endoderm specification of LRP5/6/Fz-mediated β-catenin signaling.

### Knockdown of Wnt ligands

Wnt genes of *Nematostella* are expressed in staggered domains along the OA axis (Kusserow et al., 2005; Lee et al., 2006); however, their individual roles in the OA patterning are still unclear. We showed earlier that co-expression of two Wnt genes, *Wnt1* and *Wnt3*, was sufficient to convey axial organizer capacity to any area of the *Nematostella* gastrula ectoderm, while other early Wnt ligands failed to elicit this effect (Kirillova et al., 2018; Kraus et al., 2016). However, even for *Wnt1* and *Wnt3* the possible role in axial patterning was not analyzed. In order to get some indication, which Wnt ligands might be involved in transmitting the signals patterning the *Nematostella* ectoderm along the OA axis, we analyzed the loss-of-function phenotypes of all the Wnt genes expressed in the early embryo of *Nematostella*. The following Wnt genes are active in the embryo at or prior to gastrula stage: *Wnt1, Wnt2, Wnt3, Wnt4, Wnt5, Wnt8a*, and *WntA* (Suppl. Fig. 9). RNAi of *Wnt5*, which was not very efficient with both shRNAs we used) and *WntA* did not elicit any noticeable effect on the expression of *Bra, Wnt2* and *Six3/6* in the gastrula. RNAi of the orally expressed *Wnt1* and especially *Wnt3* resulted in a reduction of the expression of the oral marker *Bra*, and its expansion to the bottom of the pharynx (Fig. 6) – a phenotype similar to the KD effect of one of the four key regulators of the oral molecular identity, *FoxB* (Lebedeva et al., 2021). RNAi of *Wnt2* and *Wnt8a*, which are normally expressed in the midbody domain, resulted in the moderate oral expansion of the aboral marker *Six3/6*, while the KD of the orally expressed *Wnt4* led to a strong aboralization of the embryo comparable to the effect of the *Fz5* KD (Fig. 6). None of the RNAi-mediated *Wnt* KDs affected *SnailA* expression (data not shown) or gastrulation.

**Figure 6.**
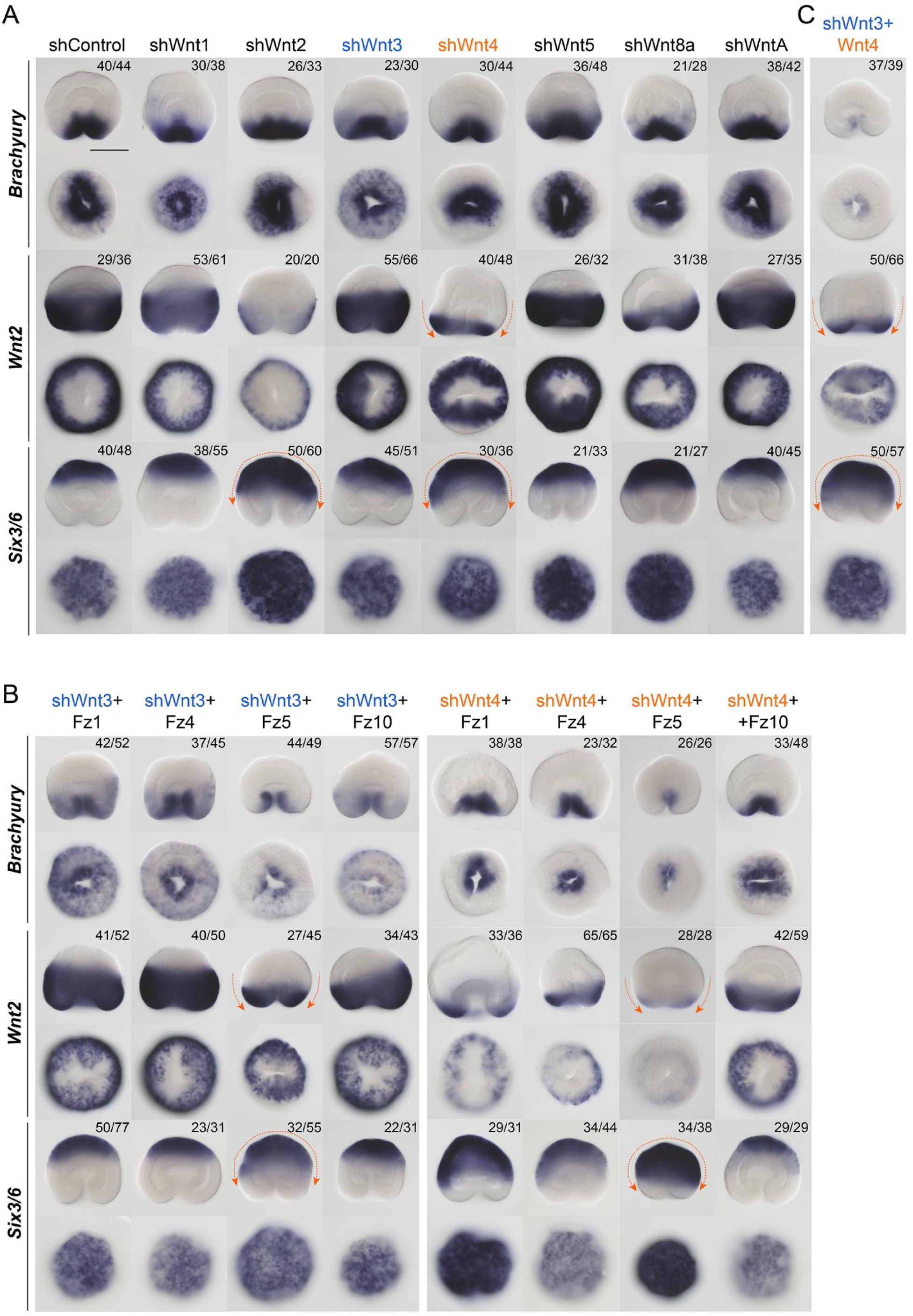
Effect of the KD of Wnt genes on the expression of the oral, midbody and aboral ectoderm markers. (A) KDs of individual *Wnt* genes. (B) Simultaneous KDs of *Wnt3* or *Wnt4* with the individual *Fz* receptor genes. (C) Simultaneous KDs of *Wnt3* and *Wnt4*. Orange arrows indicate the direction of the particularly drastic expression shifts. The numbers in the top right corner show the fraction of the embryo demonstrating this phenotype. Scale bar 100 μm. For each gene, lateral views (oral end down) on the top, oral (aboral in case of *Six3/6*) views on the bottom.

Since KDs of *Wnt3* and *Wnt4* resulted in the strongest patterning changes, we tested whether concomitant knockdowns of the individual Fz receptors would lead to a synergistic effect. Simultaneous KD of *Wnt3* with individual *Fz* receptor genes showed a more prominent reduction in the expression of the oral marker *Bra* than *Wnt3* KD alone in all combinations. However, this effect was strongest in the shWnt3+Fz10 and the shWnt3+Fz5 combinations. When shWnt4+Fz combinations were tested, the effects were even more noticeable. Oral expression of *Bra* was reduced in all Wnt4-Fz combinations in comparison to *Wnt4* RNAi. The aboralization of the embryo characteristic for the *Wnt4* KD was observed in all combinations, however, simultaneous KD of *Wnt4* and *Fz5* resulted in a more extensive aboralization than that observed upon individual KDs of *Wnt4* or *Fz5*, suggesting Fz5 as a highly probable interaction partner for Wnt4 – a hypothesis, which can be tested by biochemical analyses in the future. Finally, we tested the result of the simultaneous KD of *Wnt3* and *Wnt4*. Similar to the *LRP5/6* KD and the combined KD of all Fz receptors, shWnt3+Wnt4 resulted in a strong aboralization of the gastrula and loss of the oral structures and mesenteries by 4 dpf (Fig. 3, Fig. 6C). Thus, we conclude that Wnt3 and Wnt4 are required for the LRP5/6/Fz-dependent OA patterning and for the maintenance of the directive axis in *Nematostella*.

## Discussion

The role of Wnt/Fz-mediated signaling in development and disease is difficult to overestimate, however, the variety of signaling pathways, which may be activated by a Wnt-Fz interaction makes such investigation highly challenging. The initial hope that a multitude of vertebrate Fz receptors and a corresponding multitude of Wnt ligands would fall into an orderly system of signaling partnerships was not supported by the data. Phylogenetic analyses showed that the large diversity of the conserved Wnt gene families in Planulozoa (Cnidaria + Bilateria) is much more ancient than the Fz gene diversity found in vertebrates (Kusserow et al., 2005). Instead, non-vertebrate planulozoans normally have four Fz genes: *Fz1/2/7, Fz4, Fz5/8*, and *Fz9/10*, which have to cope with all the various Wnt ligands (Bastin et al., 2015; Janssen et al., 2015; Qian et al., 2013; Robert et al., 2014; Wijesena et al., 2022). Work on bilaterian – mostly vertebrate – models demonstrated partial redundancy of Fz receptors, as well as the involvement of the same receptors in both the cWnt, and the non-canonical Wnt signaling (Bhat, 1998; Dong et al., 2018; Fischer et al., 2007; Voloshanenko et al., 2017; Wang et al., 2016; Yu et al., 2012).

One of the Wnt-mediated signaling pathways, the cWnt or Wnt/β-catenin pathway, appears to be the oldest axial patterning system present in animals. cWnt pathway involvement in the patterning of the PA axis of Bilateria and the OA axis of Cnidaria has been convincingly demonstrated functionally during the last thirty years (Darras et al., 2018; Darras et al., 2011; Fu et al., 2012; Kiecker and Niehrs, 2001; Kraus et al., 2016; Lebedeva et al., 2021; Marlow et al., 2013; McCauley et al., 2015; Nordström et al., 2002; Prühs et al., 2017; Range et al., 2013), and expression data suggest that cWnt may also be responsible for axial patterning in the earlier branching ctenophores and sponges (Leininger et al., 2014; Pang et al., 2010). Another ancestral developmental function of β-catenin appears to be the definition of the endomesodermal and, subsequently, the endodermal domain during germ layer specification in Bilateria and Cnidaria (Henry et al., 2008; Leclère et al., 2016; Lhomond et al., 2012; Logan et al., 1999; Martín-Durán et al., 2016; Momose et al., 2008; Momose and Houliston, 2007; Wikramanayake et al., 2003). Among cnidarians, the role of Fz-mediated signaling in gastrulation and OA patterning has been addressed in a hydroid *Clytia hemisphaerica*. There, two *Fz* mRNAs, *CheFz1* (*Fz1/2/7* ortholog) and *ChFz3* (*Fz9/10* ortholog) are maternally localized to the animal and the vegetal hemispheres of the egg respectively and appear to have opposing functions. *CheFz1* KD results in a delayed endoderm formation and suppression of the animal/oral marker gene expression, while *CheFz3* KD leads to the oralization of the embryo, abolishes vegetal/aboral marker genes and accelerates the ingression of the endodermal cells (Momose and Houliston, 2007). *CheFz1* is also reported to be involved in the Strabismus/Dishevelled-mediated embryo elongation in *Clytia*, suggesting that CheFz1 is active in the cWnt as well as in the Wnt/PCP pathways (Momose et al., 2012). *CheWnt3* (*Wnt3* ortholog), whose mRNA is maternally localized to the animal pole, appears to be the key ligand responsible for the oralization, likely by signaling via CheFz1 (Momose et al., 2008). This mode of regulation, however, does not recapitulate the situation we observed in the anthozoan cnidarian model *Nematostella vectensis*. In *Nematostella, Fz1, Fz5*, and *LRP5/6* mRNAs are maternally deposited, however, these mRNAs are evenly distributed throughout the egg. Fz expression during early development is in partially overlapping domains, and it roughly recapitulates the expression of *Fz* genes in sea urchin embryos of comparable stages (Robert et al., 2014). *Nematostella Wnt* genes, with a possible exception of *Wnt5* showing some maternal transcript (Suppl. Fig. 9), as well as *Fz4* and *Fz10* are zygotically expressed. Proteomics data indicate that among the four Fz receptors, LRP5/6 and all Wnt ligands, only Fz5 protein is detectable in the *Nematostella* egg (Levitan et al., 2015). Our AZK treatment experiments suggest that β-catenin-dependent specification of the future preendodermal plate is an early event happening prior to the onset of the zygotic transcription around 6 hpf and is thus likely to rely on maternally deposited molecules. We observed normal endoderm invagination, normal expression of the endodermal markers *SnailA* and *ERG* (Fig. 4B), which are negatively controlled by β-catenin, and the lack of the expression of the β-catenin signaling targets such as *Bra* in the endoderm of the embryos treated with AZK after 6 hpf (Lebedeva et al., 2021). This indicates that after being specified, the future endoderm becomes insensitive to the modulation of the β-catenin signaling intensity. Moreover, normal endoderm specification upon knockdowns of the maternally deposited *LRP5/6, Fz1* and *Fz5* suggest that this process may not require Fz/LRP5/6-mediated signaling but rely on the cytoplasmic components of the β-catenin signaling pathway. In the future, generation and incrossing of the β-catenin^wt/-^ and LRP5/6^wt/-^ knockout lines will allow us to definitively answer the question whether or not endoderm specification relies on maternal β-catenin and is LRP5/6-independent, as our data seem to suggest at the moment. The gastrulation delay in LRP5/6 morphants also pointed out the likely involvement of LRP5/6-mediated β-catenin signaling in the process of gastrulation.

In echinoderms, the early β-catenin-dependent specification of the endomesodermal domain is followed by the segregation of the endoderm from the mesoderm, and the subsequent Wnt-dependent PA patterning. In the endoderm, β-catenin signaling remains strong, while in the mesoderm β-catenin signaling becomes suppressed (Lhomond et al., 2012; Logan et al., 1999; McCauley et al., 2015; McClay et al., 2021; Range et al., 2013; Sun et al., 2021; Wikramanayake et al., 1998; Wikramanayake et al., 2004). A similar sequence of events – the early β-catenin-dependent definition of the future endodermal domain, the formation of the boundary between the β-catenin-sensitive future oral ectoderm and the β- catenin-insensitive future endoderm, and the subsequent Wnt-dependent OA patterning of the ectoderm also occurs in *Nematostella*, and these events seem to follow the same regulatory logic as described for the sea urchin. Recently, we described the regulatory principle underlying β-catenin-dependent OA patterning of the ectoderm in *Nematostella*, which leads to the subdivision of the ectoderm into an oral, midbody, and an aboral domain (Lebedeva et al., 2021). This subdivision happens as follows: a number of transcription factor coding genes, whose expression is positively regulated by β-catenin signaling, start to be expressed in the oral hemisphere of the *Nematostella* embryo. Their expression resolves into specific domains along the oral-aboral axis because some of these genes, which are expressed more orally, encode transcriptional repressors acting on the genes, which are expressed more aborally. This creates the two main molecular boundaries of the early embryo of *Nematostella* – the oral/midbody boundary, and the midbody/aboral boundary. We showed that the oral/midbody boundary is established by the module of four transcription factors: Brachyury, Lmx, FoxA, and FoxB, while the midbody/aboral boundary is created due to the activity of the transcription factor Sp6-9 (Lebedeva et al., 2021). The whole regulatory principle and the genes involved in the OA patterning of the *Nematostella* embryo showed striking resemblance to the logic and the components of the PA patterning in deuterostome Bilateria (Darras et al., 2018; Darras et al., 2011; Kiecker and Niehrs, 2001; Lebedeva et al., 2021; Nordström et al., 2002; Range, 2018; Range et al., 2013). In contrast to endoderm specification, axial patterning is strongly affected by the knockdowns of LRP5/6 or combined knockdowns of Fz receptors, which demonstrate partial functional redundancy. The fact that LRP5/6 phenotype is phenocopied only by the simultaneous knockdown of all four Fz receptors suggests that all of them are involved in β-catenin signaling. The similarity of the combined *Wnt3*+*Wnt4* KD phenotype to the *LRP5/6* KD and the quadruple *Fz* KD indicates that these two orally expressed Wnt ligands play the main role in the Fz/LRP5/6-mediated OA patterning during early *Nematostella* development. KD phenotype similarity also suggests that among these two Wnt ligands, Wnt4 appears to be the one predominantly signalling via the aborally expressed Fz5.

Taken together, our data suggest the crucial role of the Wnt/LRP5/6/Fz-mediated signaling in the OA patterning of the sea anemone *Nematostella vectensis*, in which different Fz receptors play partially redundant roles. In contrast, we do not find evidence for the involvement of Fz/LRP5/6-mediated signaling in the specification of the preendodermal plate. With this work, we lay the foundation for the future research, which will show whether Fz functions become more distinct at later developmental stages, identify the possible signaling preferences of the different Wnt ligands towards different Fz receptors, and address the role of the non-canonical Wnt pathways in *Nematostella* development. Ultimately, it will be important to understand not only the difference between the functions of the different Fz molecules but also the role of their redundancy and the selective pressures maintaining what appears to be an ancestral Fz redundancy conserved in Cnidaria and Bilateria.

## Materials and Methods

### Animals, microinjection and electroporation

Adult *Nematostella vectensis* polyps were kept separated by sex in 16‰ artificial sea water (*Nematostella* medium = NM) at 18°C in the dark. Spawning was induced by placing the polyps into an illuminated incubator set to 25°C for 10 hrs. The eggs were de-jellied with 3% L-cystein/NM as described in (Genikhovich and Technau, 2009). Microinjection of the shRNAs and morpholinos and electroporation of shRNAs against maternally expressed transcripts was performed prior to fertilization. For zygotic transcripts, electroporation and microinjection was performed after fertilization. The embryos were raised at 21°C.

### Gene knockdown, mRNA overexpression and inhibitor treatments

shRNA-mediated gene knockdown was performed as described in (Karabulut et al., 2019). Two independent, non-overlapping shRNAs were used for each gene to make sure that the KD result was specific. RNAi efficiency was tested by in situ hybridization (Suppl. Fig. 2A-B). For morpholino KDs, the activity of the morpholinos was confirmed by co-injecting each of them with *mCherry* mRNA containing the recognition sequence for the respective morpholino oligonucleotide and testing whether mCherry translation was suppressed in comparison to the situation, when the same mRNA was co-injected with a control MO (Suppl. Fig. 2C). mRNA was synthesized with mMessage mMachine kit (Life Technologies) and purified with the Monarch RNA clean-up kit (NEB). 5μM 1-azakenpaullone (Sigma) used for the treatments was prepared by diluting 5 mM AZK dissolved in DMSO with NM. Equal volume of DMSO was used to treat the control embryos. The duration of the treatment is described on Fig. 4A. The recognition sequences for the shRNAs and morpholino sequences are shown in the Supplementary Tables 1 and 2. Accession numbers for the genes used in the study are presented in the Supplementary Table 3.

### In situ hybridization and phalloidin staining

In situ hybridization was performed as described in (Kraus et al., 2016) with a single change: the embryos were fixed for 1 hour at room temperature in 4%PFA/PBS, washed several times in PTw (1× PBS, 0.1% Tween 20), then in 100% methanol, and finally stored at −20 °C. Dig-labelled RNA probes were detected with anti-Digoxigenin-AP Fab fragments (Roche) diluted 1:4000 in 0.5% blocking reagent (Roche) in 1× MAB. After unbound antibody was removed by a series of ten 10-minute PTw washes, the embryos were stained with a mixture of NBT/BCIP, embedded in 86% glycerol and imaged using a Nikon 80i compound microscope equipped with the Nikon DS-Fi1 camera. For phalloidin staining, the embryos were fixed in 4%PFA/PTwTx (1× PBS, 0.1% Tween 20, 0.2% Triton X100) for 1 h at room temperature, washed 5 times with PTwTx, incubated in 100% acetone pre-cooled to −20 °C for 7 min on ice, and washed 3 more times with PTwTx. 2 μL of phalloidin-AlexaFluor488 (ThermoFisher) was added per 100 μL PTwTx, and the embryos were stained overnight at 4°C. After eight 10-minute washes with PTwTx, the embryos were gradually embedded in Vectashield (Vectorlabs) and imaged with the Leica SP8 CLSM.

## Acknowledgements

This work has been funded by the Austrian Science Foundation (FWF) grant P30404-B29 to G.G.. We are grateful to the Core Facility for Cell Imaging and Ultrastructure Research of the University of Vienna for the access to the confocal microscope. We thank David Mörsdorf for his valuable comments on the manuscript.

## Author contributions

I.N. performed experiments, analyzed data and wrote the paper, T.L. performed experiments and edited the paper, G.G. conceived the study, performed experiments, analyzed data and wrote the paper.

## Data availability

All data needed to evaluate the conclusions in the paper are present in the paper or the supplementary materials.

## References

Aberle, H., Bauer, A., Stappert, J., Kispert, A. and Kemler, R. (1997). beta-catenin is a target for the ubiquitin-proteasome pathway. EMBO J 16, 3797–3804.

Acebron, S. P. and Niehrs, C. (2016). beta-Catenin-Independent Roles of Wnt/LRP6 Signaling. Trends Cell Biol 26, 956–967.

Adamska, M., Larroux, C., Adamski, M., Green, K., Lovas, E., Koop, D., Richards, G. S., Zwafink, C. and Degnan, B. M. (2010). Structure and expression of conserved Wnt pathway components in the demosponge Amphimedon queenslandica. Evol Dev 12, 494–518.

Bastin, B. R., Chou, H. C., Pruitt, M. M. and Schneider, S. Q. (2015). Structure, phylogeny, and expression of the frizzled-related gene family in the lophotrochozoan annelid Platynereis dumerilii. Evodevo 6, 37.

Bhat, K. M. (1998). frizzled and frizzled 2 play a partially redundant role in wingless signaling and have similar requirements to wingless in neurogenesis. Cell 95, 1027–1036.

Croce, J. C. and McClay, D. R. (2008). Evolution of the Wnt pathways. Methods Mol Biol 469, 3–18.

Darras, S., Fritzenwanker, J. H., Uhlinger, K. R., Farrelly, E., Pani, A. M., Hurley, I. A., Norris, R. P., Osovitz, M., Terasaki, M., Wu, M., et al. (2018). Anteroposterior axis patterning by early canonical Wnt signaling during hemichordate development. PLoS Biol 16, e2003698.

Darras, S., Gerhart, J., Terasaki, M., Kirschner, M. and Lowe, C. J. (2011). beta-catenin specifies the endomesoderm and defines the posterior organizer of the hemichordate Saccoglossus kowalevskii. Development 138, 959–970.

Dong, B., Vold, S., Olvera-Jaramillo, C. and Chang, H. (2018). Functional redundancy of frizzled 3 and frizzled 6 in planar cell polarity control of mouse hair follicles. Development 145.

Fischer, T., Guimera, J., Wurst, W. and Prakash, N. (2007). Distinct but redundant expression of the Frizzled Wnt receptor genes at signaling centers of the developing mouse brain. Neuroscience 147, 693–711.

Flack, J. E., Mieszczanek, J., Novcic, N. and Bienz, M. (2017). Wnt-Dependent Inactivation of the Groucho/TLE Co-repressor by the HECT E3 Ubiquitin Ligase Hyd/UBR5. Mol Cell 67, 181–193 e185.

Fu, J., Posnien, N., Bolognesi, R., Fischer, T. D., Rayl, P., Oberhofer, G., Kitzmann, P., Brown, S. J. and Bucher, G. (2012). Asymmetrically expressed axin required for anterior development in Tribolium. Proc Natl Acad Sci USA 109, 7782–7786.

Gammons, M. and Bienz, M. (2018). Multiprotein complexes governing Wnt signal transduction. Curr Opin Cell Biol 51, 42–49.

Garcia de Herreros, A. and Dunach, M. (2019). Intracellular Signals Activated by Canonical Wnt Ligands Independent of GSK3 Inhibition and beta-Catenin Stabilization. Cells 8.

Genikhovich, G., Fried, P., Prünster, M. M., Schinko, J. B., Gilles, A. F., Meier, K., Iber, D. and Technau, U. (2015). Axis patterning by BMPs: cnidarian network reveals evolutionary constraints. Cell Rep 10, 1646–1654.

Genikhovich, G. and Technau, U. (2009). Induction of spawning in the starlet sea anemone Nematostella vectensis, in vitro fertilization of gametes, and dejellying of zygotes. CSH protocols 2009, pdb prot5281.

Grainger, S. and Willert, K. (2018). Mechanisms of Wnt signaling and control. Wiley Interdiscip Rev Syst Biol Med, e1422.

Helm, R. R., Siebert, S., Tulin, S., Smith, J. and Dunn, C. W. (2013). Characterization of differential transcript abundance through time during Nematostella vectensis development. BMC Genomics 14, 266.

Henry, J. Q., Perry, K. J., Wever, J., Seaver, E. and Martindale, M. Q. (2008). Beta-catenin is required for the establishment of vegetal embryonic fates in the nemertean, Cerebratulus lacteus. Dev Biol 317, 368–379.

Janssen, R., Schönauer, A., Weber, M., Turetzek, N., Hogvall, M., Goss, G., Patel, N., McGregor, A. and Hilbrant, M. (2015). The evolution and expression of panarthropod frizzled genes. Frontiers Ecol Evol 3, 96.

Karabulut, A., He, S., Chen, C. Y., McKinney, S. A. and Gibson, M. C. (2019). Electroporation of short hairpin RNAs for rapid and efficient gene knockdown in the starlet sea anemone, Nematostella vectensis. Dev Biol 448, 7–15.

Kiecker, C. and Niehrs, C. (2001). A morphogen gradient of Wnt/beta-catenin signalling regulates anteroposterior neural patterning in Xenopus. Development 128, 4189–4201.

Kirillova, A., Genikhovich, G., Pukhlyakova, E., Demilly, A., Kraus, Y. and Technau, U. (2018). Germ-layer commitment and axis formation in sea anemone embryonic cell aggregates. Proc Natl Acad Sci USA 115, 1813–1818.

Kraus, Y., Aman, A., Technau, U. and Genikhovich, G. (2016). Pre-bilaterian origin of the blastoporal axial organizer. Nat Commun 7, 11694.

Kusserow, A., Pang, K., Sturm, C., Hrouda, M., Lentfer, J., Schmidt, H. A., Technau, U., von Haeseler, A., Hobmayer, B., Martindale, M. Q., et al. (2005). Unexpected complexity of the Wnt gene family in a sea anemone. Nature 433, 156–160.

Lebedeva, T., Aman, A. J., Graf, T., Niedermoser, I., Zimmermann, B., Kraus, Y., Schatka, M., Demilly, A., Technau, U. and Genikhovich, G. (2021). Cnidarian-bilaterian comparison reveals the ancestral regulatory logic of the β-catenin dependent axial patterning. Nat Commun 12, 4032.

Leclère, L., Bause, M., Sinigaglia, C., Steger, J. and Rentzsch, F. (2016). Development of the aboral domain in Nematostella requires beta-catenin and the opposing activities of Six3/6 and Frizzled5/8. Development 143, 1766–1777.

Leclère, L. and Rentzsch, F. (2014). RGM Regulates BMP-Mediated Secondary Axis Formation in the Sea Anemone Nematostella vectensis. Cell Rep 9, 1–10.

Lee, P. N., Pang, K., Matus, D. Q. and Martindale, M. Q. (2006). A WNT of things to come: evolution of Wnt signaling and polarity in cnidarians. Seminars Cell Dev Biol 17, 157–167.

Leininger, S., Adamski, M., Bergum, B., Guder, C., Liu, J., Laplante, M., Brate, J., Hoffmann, F., Fortunato, S., Jordal, S., et al. (2014). Developmental gene expression provides clues to relationships between sponge and eumetazoan body plans. Nat Commun 5, 3905.

Levitan, S., Sher, N., Brekhman, V., Ziv, T., Lubzens, E. and Lotan, T. (2015). The making of an embryo in a basal metazoan: Proteomic analysis in the sea anemone Nematostella vectensis. Proteomics 15, 4096–4104.

Lhomond, G., McClay, D. R., Gache, C. and Croce, J. C. (2012). Frizzled1/2/7 signaling directs beta-catenin nuclearisation and initiates endoderm specification in macromeres during sea urchin embryogenesis. Development 139, 816–825.

Logan, C. Y., Miller, J. R., Ferkowicz, M. J. and McClay, D. R. (1999). Nuclear beta-catenin is required to specify vegetal cell fates in the sea urchin embryo. Development 126, 345–357.

MacDonald, B. T. and He, X. (2012). Frizzled and LRP5/6 receptors for Wnt/beta-catenin signaling. Cold Spring Harb Perspect Biol 4.

Marlow, H., Matus, D. Q. and Martindale, M. Q. (2013). Ectopic activation of the canonical wnt signaling pathway affects ectodermal patterning along the primary axis during larval development in the anthozoan Nematostella vectensis. Dev Biol 380, 324–334.

Martín-Durán, J. M., Passamaneck, Y. J., Martindale, M. Q. and Hejnol, A. (2016). The developmental basis for the recurrent evolution of deuterostomy and protostomy. Nature Ecol. Evol. 1, 5.

McCauley, B. S., Akyar, E., Saad, H. R. and Hinman, V. F. (2015). Dose-dependent nuclear beta-catenin response segregates endomesoderm along the sea star primary axis. Development 142, 207–217.

McClay, D. R., Croce, J. C. and Warner, J. F. (2021). Conditional specification of endomesoderm. Cells Dev 167, 203716.

Momose, T., Derelle, R. and Houliston, E. (2008). A maternally localised Wnt ligand required for axial patterning in the cnidarian Clytia hemisphaerica. Development 135, 2105–2113.

Momose, T. and Houliston, E. (2007). Two oppositely localised frizzled RNAs as axis determinants in a cnidarian embryo. PLoS Biol 5, e70.

Momose, T., Kraus, Y. and Houliston, E. (2012). A conserved function for Strabismus in establishing planar cell polarity in the ciliated ectoderm during cnidarian larval development. Development 139, 4374–4382.

Nordström, U., Jessell, T. M. and Edlund, T. (2002). Progressive induction of caudal neural character by graded Wnt signaling. Nature Neurosci 5, 525–532.

Pang, K., Ryan, J. F., Mullikin, J. C., Baxevanis, A. D. and Martindale, M. Q. (2010). Genomic insights into Wnt signaling in an early diverging metazoan, the ctenophore Mnemiopsis leidyi. Evodevo 1, 10.

Park, H. W., Kim, Y. C., Yu, B., Moroishi, T., Mo, J. S., Plouffe, S. W., Meng, Z., Lin, K. C., Yu, F. X., Alexander, C. M., et al. (2015). Alternative Wnt Signaling Activates YAP/TAZ. Cell 162, 780–794.

Prühs, R., Beermann, A. and Schroder, R. (2017). The Roles of the Wnt-Antagonists Axin and Lrp4 during Embryogenesis of the Red Flour Beetle Tribolium castaneum. J Dev Biol 5, 10.

Qian, G., Li, G., Chen, X. and Wang, Y. (2013). Characterization and embryonic expression of four amphioxus Frizzled genes with important functions during early embryogenesis. Gene Expr Patterns 13, 445–453.

Range, R. C. (2018). Canonical and non-canonical Wnt signaling pathways define the expression domains of Frizzled 5/8 and Frizzled 1/2/7 along the early anterior-posterior axis in sea urchin embryos. Dev Biol 444, 83–92.

Range, R. C., Angerer, R. C. and Angerer, L. M. (2013). Integration of Canonical and Noncanonical Wnt Signaling Pathways Patterns the Neuroectoderm Along the Anterior–Posterior Axis of Sea Urchin Embryos. PLoS biology 11, e1001467.

Robert, N., Lhomond, G., Schubert, M. and Croce, J. C. (2014). A comprehensive survey of wnt and frizzled expression in the sea urchin Paracentrotus lividus. Genesis 52, 235–250.

Röttinger, E., Dahlin, P. and Martindale, M. Q. (2012). A Framework for the Establishment of a Cnidarian Gene Regulatory Network for “Endomesoderm” Specification: The Inputs of ss-Catenin/TCF Signaling. PLoS Gen 8, e1003164.

Saina, M., Genikhovich, G., Renfer, E. and Technau, U. (2009). BMPs and chordin regulate patterning of the directive axis in a sea anemone. Proc Natl Acad Sci USA 106, 18592–18597.

Schenkelaars, Q., Fierro-Constain, L., Renard, E., Hill, A. L. and Borchiellini, C. (2015). Insights into Frizzled evolution and new perspectives. Evol Dev 17, 160–169.

Semenov, M. V., Habas, R., Macdonald, B. T. and He, X. (2007). SnapShot: Noncanonical Wnt Signaling Pathways. Cell 131, 1378.

Sun, H., Peng, C. J., Wang, L., Feng, H. and Wikramanayake, A. H. (2021). An early global role for Axin is required for correct patterning of the anterior-posterior axis in the sea urchin embryo. Development 148, dev191197.

Tamai, K., Zeng, X., Liu, C., Zhang, X., Harada, Y., Chang, Z. and He, X. (2004). A mechanism for Wnt coreceptor activation. Molecular cell 13, 149–156.

van Amerongen, R. and Nusse, R. (2009). Towards an integrated view of Wnt signaling in development. Development 136, 3205–3214.

Villarroel, A., Del Valle-Perez, B., Fuertes, G., Curto, J., Ontiveros, N., Garcia de Herreros, A. and Dunach, M. (2020). Src and Fyn define a new signaling cascade activated by canonical and non-canonical Wnt ligands and required for gene transcription and cell invasion. Cell Mol Life Sci 77, 919–935.

Voloshanenko, O., Gmach, P., Winter, J., Kranz, D. and Boutros, M. (2017). Mapping of Wnt-Frizzled interactions by multiplex CRISPR targeting of receptor gene families. FASEB J 31, 4832–4844.

Wang, Y., Chang, H., Rattner, A. and Nathans, J. (2016). Frizzled Receptors in Development and Disease. Curr Top Dev Biol 117, 113–139.

Warner, J. F., Guerlais, V., Amiel, A. R., Johnston, H., Nedoncelle, K. and Rottinger, E. (2018). NvERTx: a gene expression database to compare embryogenesis and regeneration in the sea anemone Nematostella vectensis. Development 145, dev162867.

Wijesena, N., Sun, H., Kumburegama, S. and Wikramanayake, A. H. (2022). Distinct Frizzled receptors independently mediate endomesoderm specification and primary archenteron invagination during gastrulation in Nematostella. Dev Biol 481, 215–225.

Wikramanayake, A. H., Hong, M., Lee, P. N., Pang, K., Byrum, C. A., Bince, J. M., Xu, R. and Martindale, M. Q. (2003). An ancient role for nuclear beta-catenin in the evolution of axial polarity and germ layer segregation. Nature 426, 446–450.

Wikramanayake, A. H., Huang, L. and Klein, W. H. (1998). beta-Catenin is essential for patterning the maternally specified animal-vegetal axis in the sea urchin embryo. Proc Natl Acad Sci USA 95, 9343–9348.

Wikramanayake, A. H., Peterson, R., Chen, J., Huang, L., Bince, J. M., McClay, D. R. and Klein, W. H. (2004). Nuclear beta-catenin-dependent Wnt8 signaling in vegetal cells of the early sea urchin embryo regulates gastrulation and differentiation of endoderm and mesodermal cell lineages. Genesis 39, 194–205.

Willert, K., Shibamoto, S. and Nusse, R. (1999). Wnt-induced dephosphorylation of axin releases beta-catenin from the axin complex. Genes Dev 13, 1768–1773.

Yu, H., Ye, X., Guo, N. and Nathans, J. (2012). Frizzled 2 and frizzled 7 function redundantly in convergent extension and closure of the ventricular septum and palate: evidence for a network of interacting genes. Development 139, 4383–4394.

