## Supplementary Information for "Sea anemone Frizzled receptors play partially redundant roles in the oral-aboral axis patterning"

### Contents

Supplementary Figures 1-8

Supplementary Tables 1-3

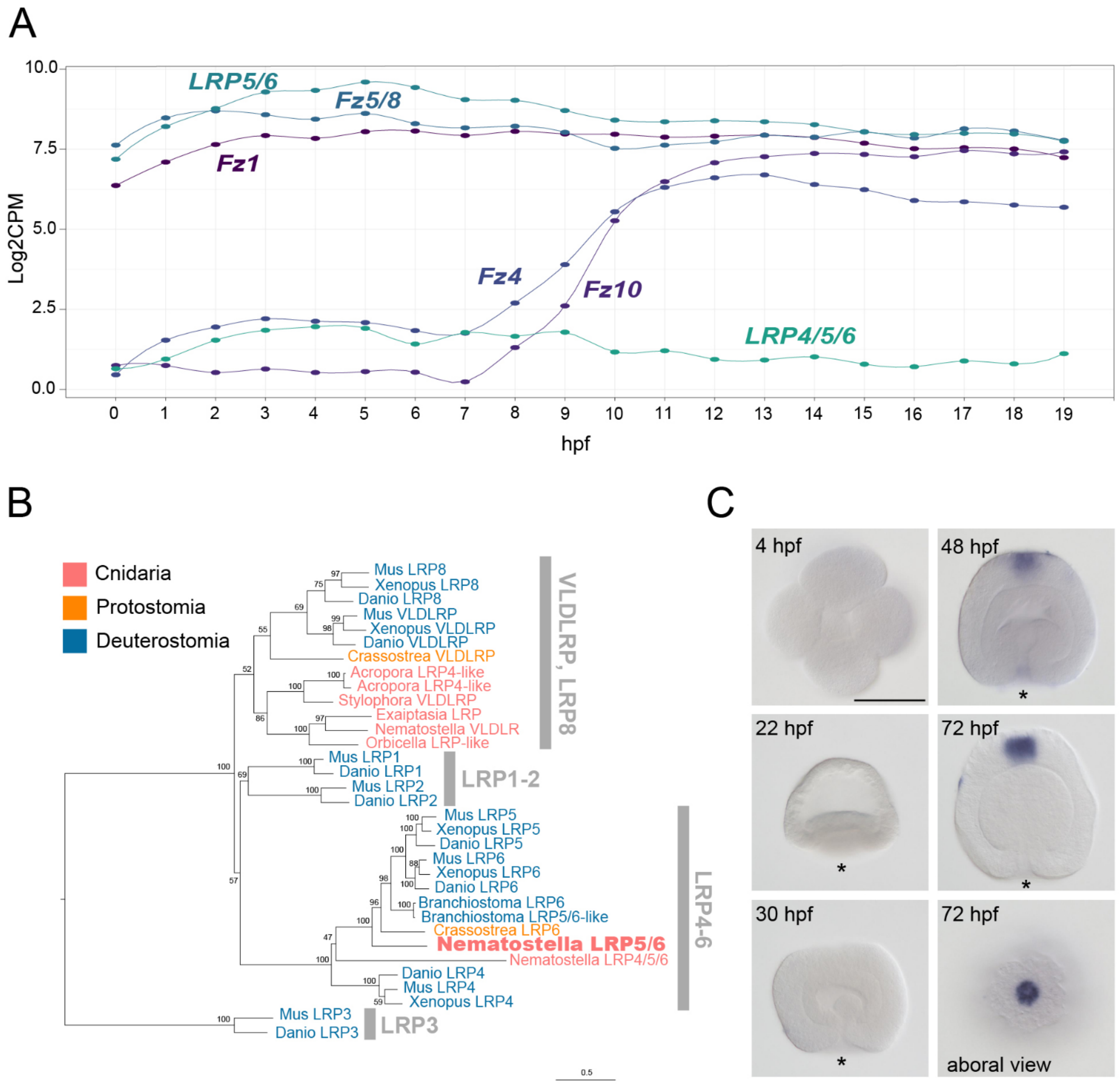


**Supplementary Figure 1.** (A) Expression dynamics of the *LRP5/6*, *LRP4/5/6-like* and *Fz* genes in the first 19 hours of *Nematostella* development according to the NvERTx database (Helm et al., 2013; Warner et al., 2018). (B) Maximum likelihood phylogeny of the LRP proteins (WAG+G4, bootstrap 100). (C) LRP4/5/6-like is expressed in the apical organ of the planula. Scale bar 100 µm.

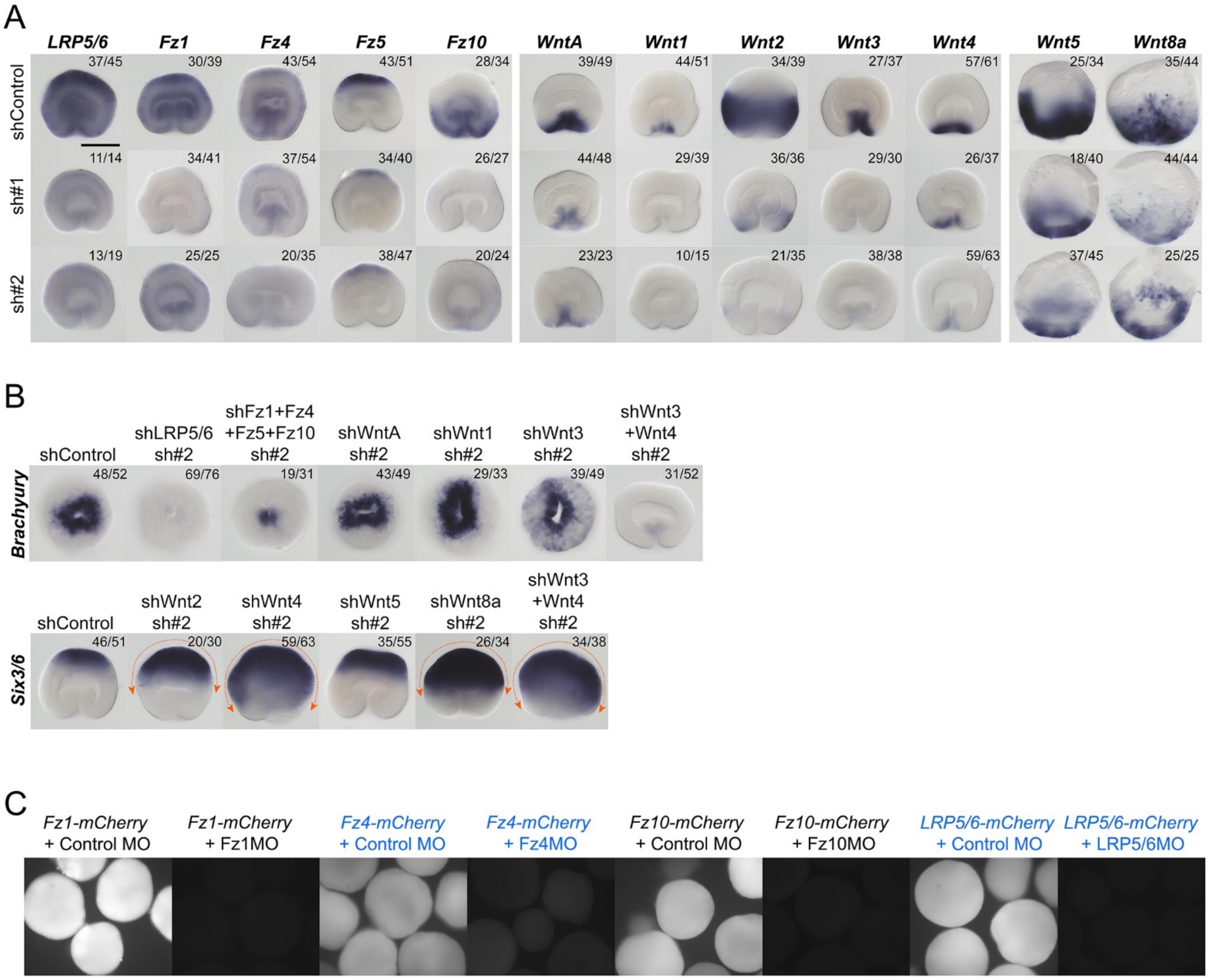


**Supplementary Figure 2.** Controls of the efficiency of the shRNAs and morpholino oligonucleotides. (A) Efficiency of the first (sh#1) and the second (sh#2) shRNA for each gene analyzed by in situ hybridization. Lateral views are shown. Wnt5 and Wnt8a expression has been assessed at mid-blastula stage because unlike all the other genes shown on the figure, Wnt5 and Wnt8a are nearly not expressed at late gastrula stage. (B) The phenotypes obtained with the sh#1 (shown on all other figures) are reproduced with minimal differences using the sh#2. Oral view is shown for Bra, lateral view is shown for Six3/6. On (A) and (B), the numbers in the top right corner show the fraction of the embryo demonstrating this phenotype. Scale bar 100 µm. (C) In vivo fluorescence shows that *mCherry* mRNA carrying the morpholino recognition sequence for the Fz1MO, Fz4MO, Fz10MO or LRP5/6MO is efficiently translated when co-injected into zygotes together with the control morpholino, but not when co-injected with the morpholinos against Fz1, Fz4, Fz10 or LRP5/6MO, respectively.

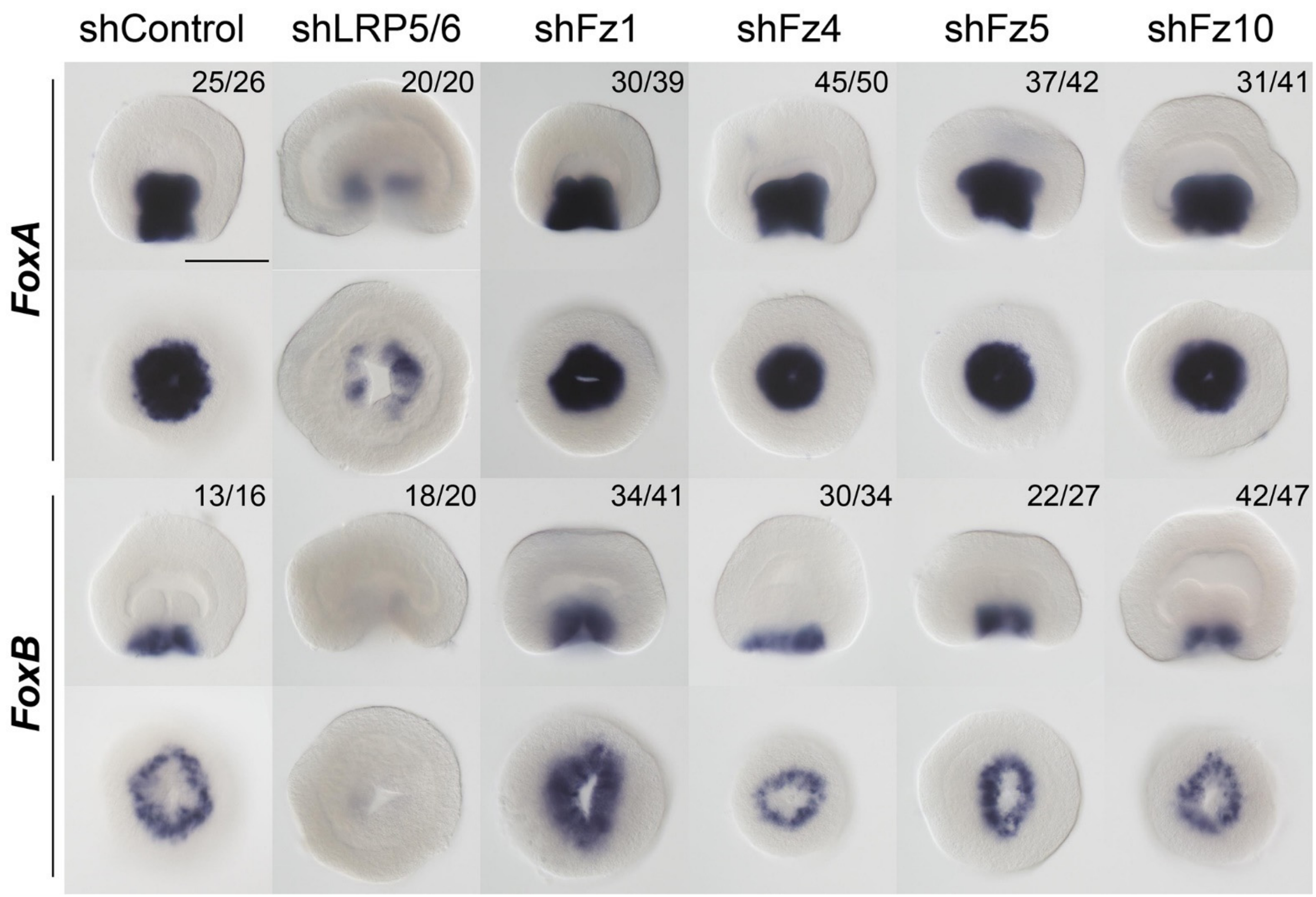


**Supplementary Figure 3.** Expression of the oral markers *FoxA* and *FoxB* upon KDs of *LRP5/6* and individual *Fz*. The numbers in the top right corner show the fraction of the embryo demonstrating this phenotype. For each gene, lateral views (oral end down) on the top, oral views on the bottom. Scale bar 100 µm.

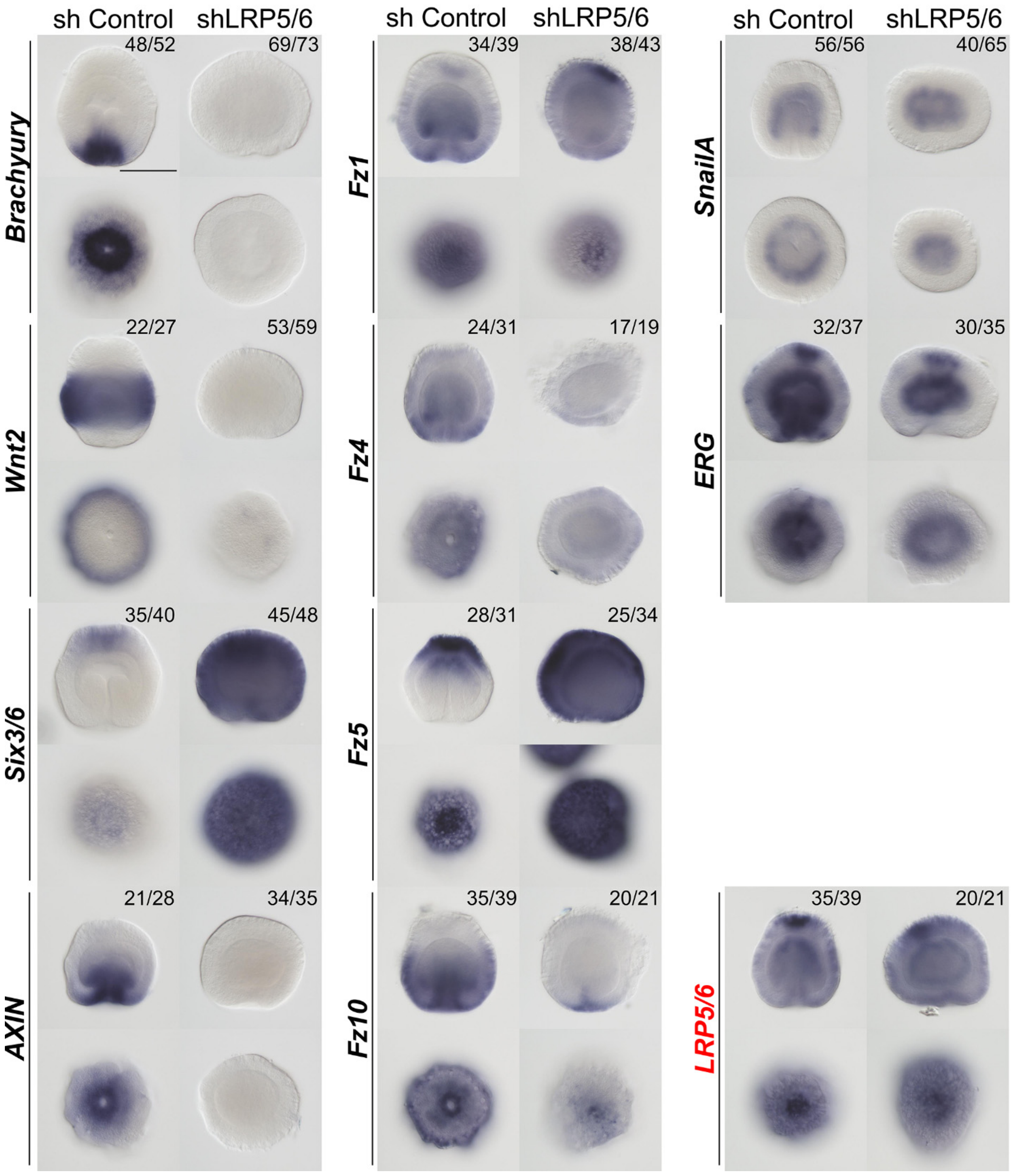


**Supplementary Figure 4.** Marker gene expression in 3 dpf planulae upon *LRP5/6* RNAi. The numbers in the top right corner show the fraction of the embryo demonstrating this phenotype. Scale bar 100 µm. For each gene, lateral views (oral end down) on the top, oral views on the bottom.

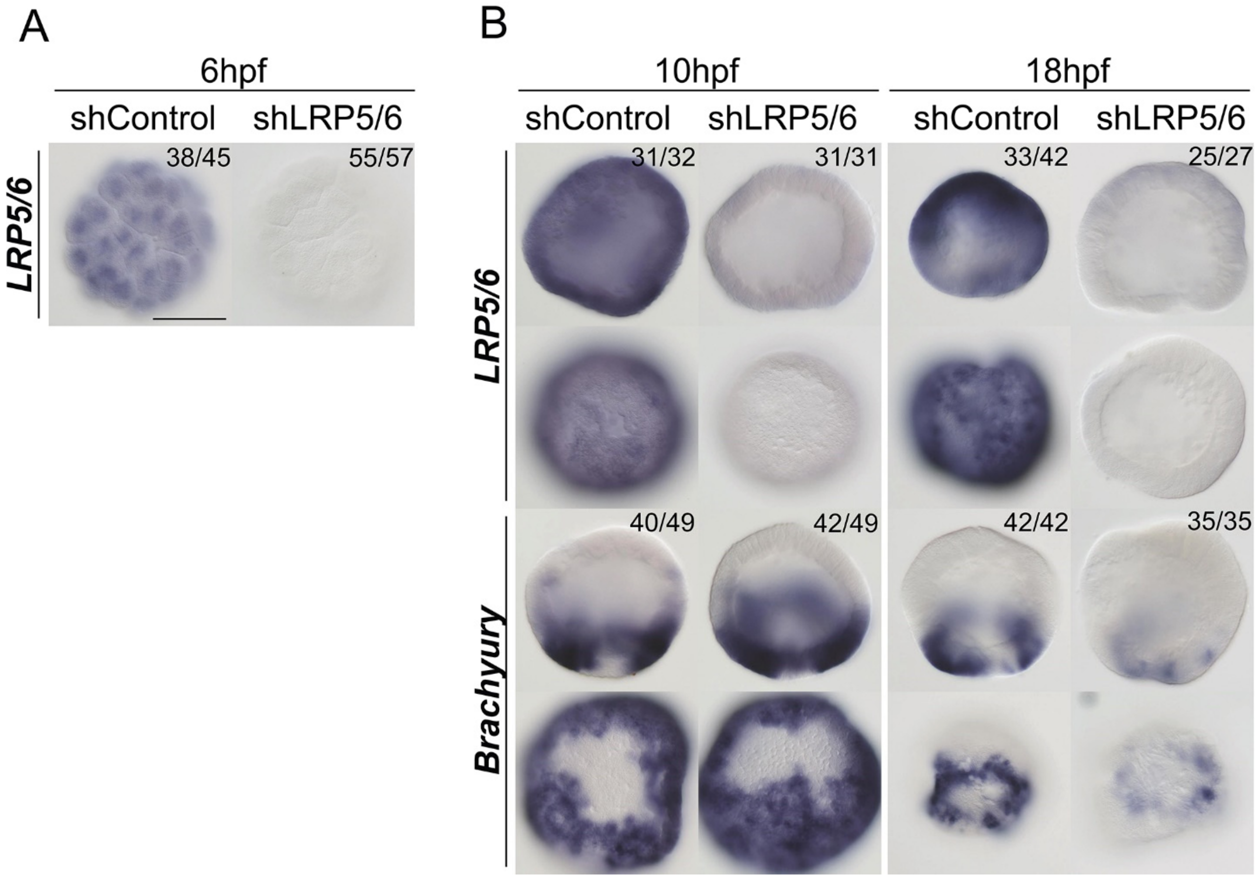


**Supplementary Figure 5.** The onset of the effect of the *LRP5/6* RNAi. (A) *LRP5/6* expression is abolished at 6 hpf. (B) *Bra* expression is not affected at 10 hpf, but starts to be suppressed by 18 hpf. The numbers in the top right corner show the fraction of the embryo demonstrating this phenotype. Scale bar 100 µm. For each gene, lateral views (oral end down) on the top, oral views on the bottom.

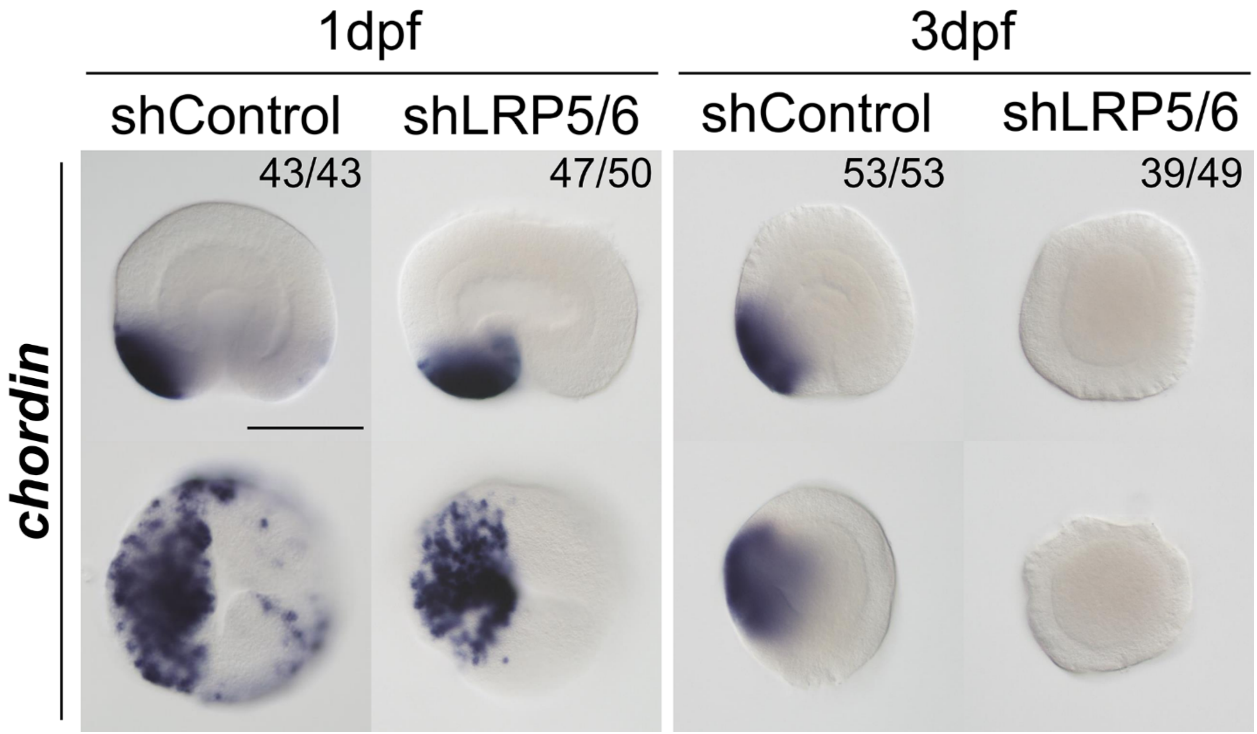


**Supplementary Figure 6.** At late gastrula (1 dpf), asymmetric *Chordin* expression indicates the establishment of the directive axis, which disappears by mid-planula (3 dpf). The numbers in the top right corner show the fraction of the embryo demonstrating this phenotype. Scale bar 100 µm. Lateral views (oral end down) on the top, oral views on the bottom.

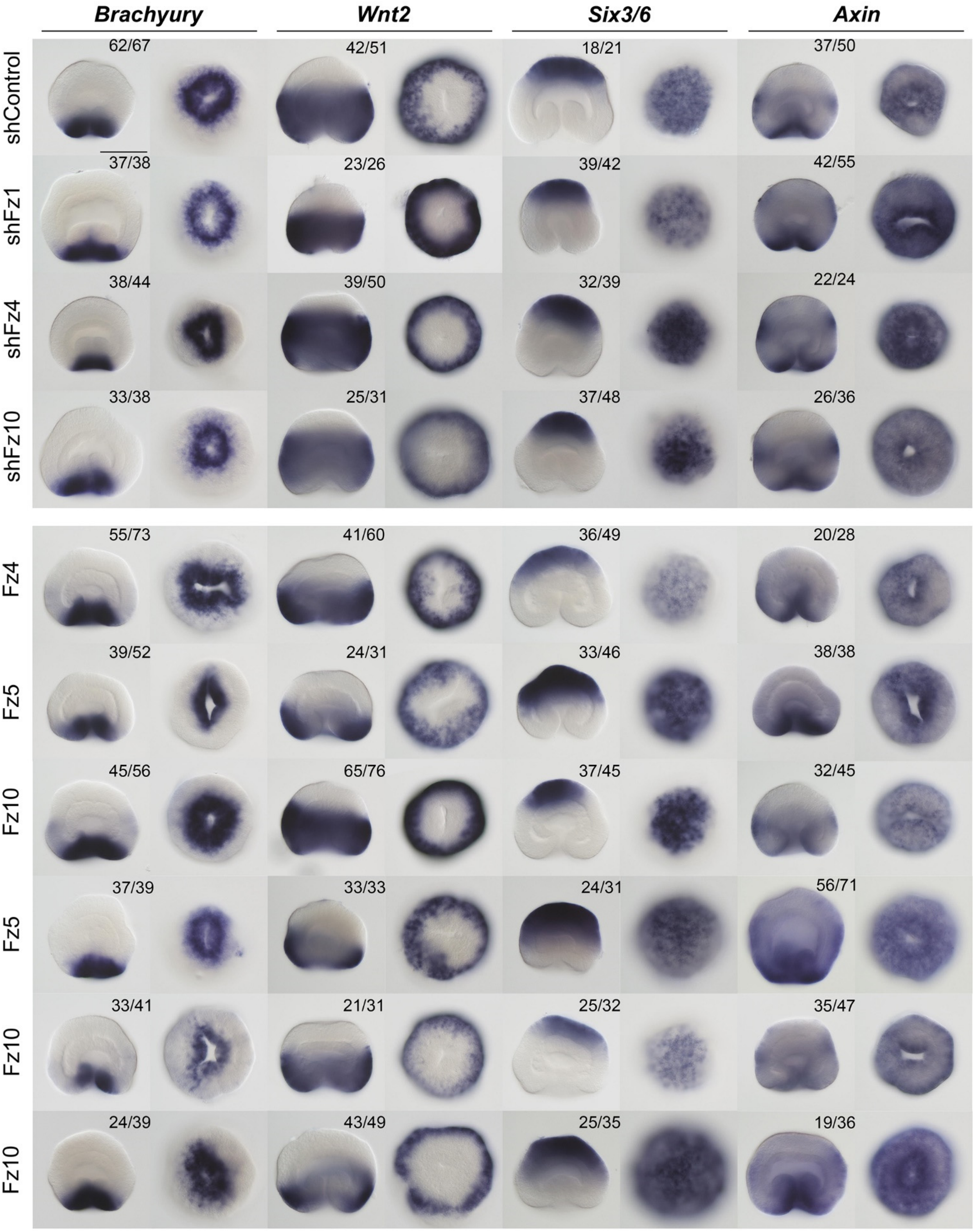


**Supplementary Figure 7.** Effects of the individual RNAi of the orally expressed Fz genes and effects of the simultaneous RNAi of all possible combinations of two Fz genes on the expression of the β-catenin-dependent markers of different axial domains. The numbers in the top right corner show the fraction of the embryo demonstrating this phenotype. Scale bar 100 µm. For each gene, lateral views (oral end down) on the left, oral (aboral in case of *Six3/6*) views on the right.

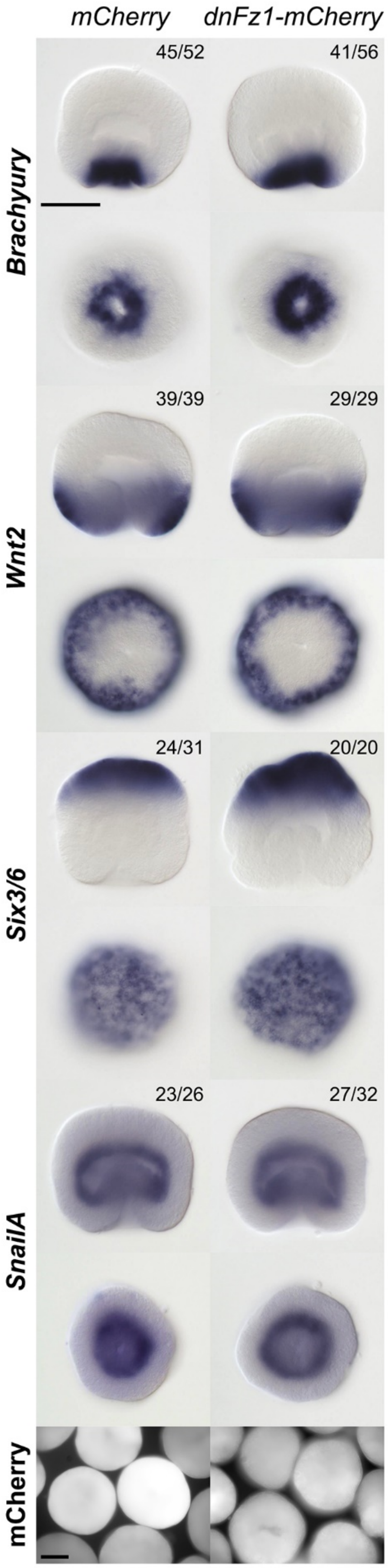


**Supplementary Figure 8.** Microinjection of the *mCherry* mRNA and *dnFz-mCherry* mRNA show no effect on the expression of the markers of the distinct axial domains in the ectoderm, and on the endodermal marker *SnailA*. The numbers in the top right corner show the fraction of the embryo demonstrating this phenotype. For each gene, lateral views (oral end down) on the top, oral (aboral in case of *Six3/6*) views on the bottom. Lower panel – mCherry fluorescence of the microinjected embryos. Scale bars 100 µm.

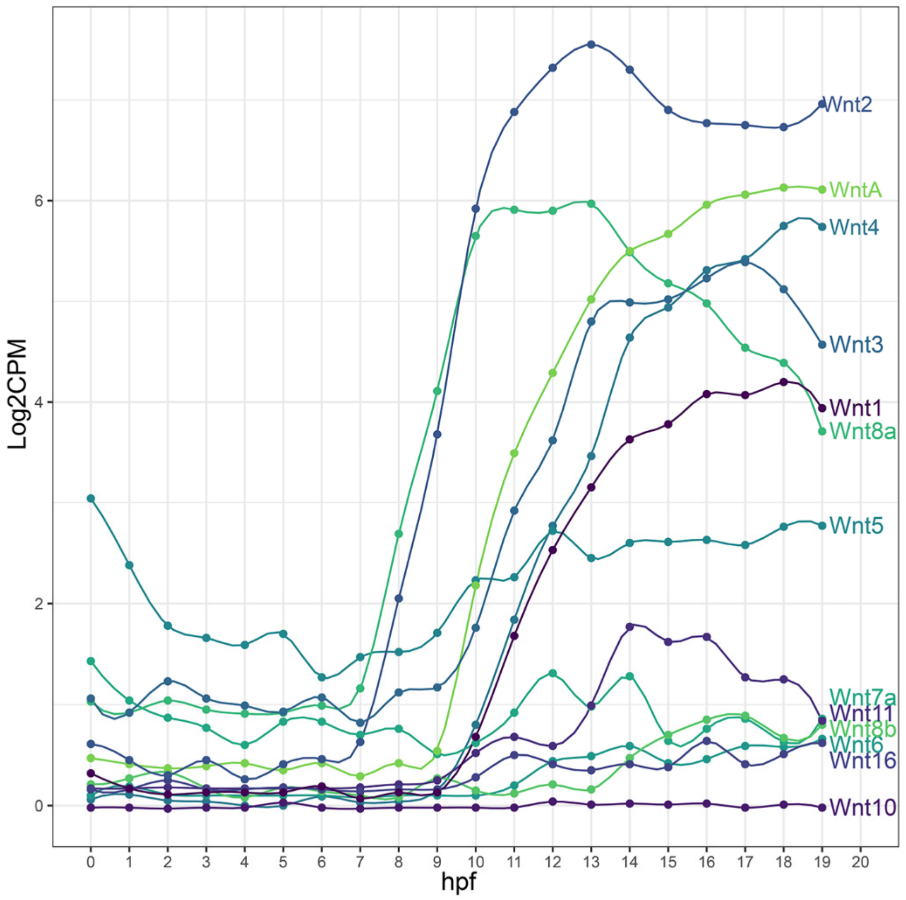


**Supplementary Figure 9.** The dynamics of the *Wnt* gene expression in the first 19 hours of *Nematostella* development according to the NvERTx database (Helm et al., 2013; Warner et al., 2018).

**Supplementary Table 1. Target sequences of the shRNAs**

| | |
|---|---|
| shLRP5/6 sh#1 | GAGAGCCTTCCACTTGTAA |
| shLRP5/6 sh#2 | GAGGAATCGTCGCATCTAT |
| shFz1 sh#1 | GAAGGCTGCACGGTTATTT |
| shFz1sh#2 | GCTTGCAATGAGCCTATCA |
| shFz4 sh#1 | GTTCAAAGCACCGAGTCTT |
| shFz4sh#2 | GCCTGAGAAACCTAGACCA |
| shFz5 sh#1 | GCGGAATAGGCTACAATTT |
| shFz5 sh#2 | GCCGGAATGAAATGGTCAA |
| shFz10 sh#1 | GGATGAACTGACAGGTCTT |
| shFz10 sh#2 | GGACAGTACCAGCAATACA |
| shWntA sh#1 | GGATAACATGGGCAAGACA |
| shWntA sh#2 | GGCGTACTATGCCAAACTT |
| shWnt1 sh#1 | GGAGGATGCAGTGATAACA |
| shWnt1 sh#2 | GGGATTTCCGTGCTCAGAT |
| shWnt2 sh#1 | GAGGGCGTTGATGAACTTA |
| shWnt2 sh#2 | GAGGATTCGCCCAATTACT |
| shWnt3 sh#1 | GGAAGACAGTGCAACTACA |
| shWnt3 sh#2 | GAGACCTCACCAAACTACT |
| shWnt4 sh#1 | GCTTCGCTAGTGTACTCAA |
| shWnt4 sh#2 | GAAATTCGATGGAGCTACT |
| shWnt5 sh#1 | GGTGCCGATGCAAGTTTCA |
| shWnt5 sh#2 | GCTCGGACTCTTATGAACT |
| shWnt8a sh#1 | GGCGCAAAGCTGTTAAGAA |
| shWnt8a sh#2 | GCAGCCTGGTCTTCCTAAA |

**Supplementary Table 2. Morpholino sequences**

| | |
|---|---|
| Control MO | GATGTGCCTAGGGTACAACAACAAT |
| Fz1MO | GCATAATCCCGGCGATTAAACTACG |
| Fz4MO | GTGACATTTTGCACGAATGGAGAAC |
| Fz10MO | AAGCTAAACGCTTAGCCCCCATATC |
| LRP5/6MO | ACAAAACAACTTTGGCGAACATCCT |

**Supplementary Table 3. Genbank accession numbers**

| Gene name | Accession |
|---|---|
| *Axin* | XP_001640692 |
| *Brachyury* | XP_032233913 |
| *Chordin* | XP_001633548 |
| *ERG* | XP_032236866 |
| *FoxA* | XP_001634555 |
| *FoxB* | XP_001631625 |
| *Fz1* | XP_001647540 |
| *Fz10* | XP_032235151 |
| *Fz4* | XP_001622965 |
| *Fz5* | XP_001634995 |
| *LRP4/5/6-like* | XM_032365558 |
| *LRP5/6* | XP_032222612 |
| *Six3/6* | XP_032228424 |
| *SnailA* | XP_032243077 |
| *Wnt1* | XP_001641494 |
| *Wnt2* | XP_032238966 |
| *Wnt3* | XP_032241388 |
| *Wnt4* | XP_001623100 |
| *Wnt5* | XP_001630693 |
| *Wnt8a* | XP_001630032 |
| *WntA* | XP_001637670 |
